# Monoclonal Antibody Developability from Early Assay Panels: Machine Learning for Formulation and Pharmacokinetic Risk

**DOI:** 10.1101/2025.10.31.685722

**Authors:** Sabitoj Singh Virk, Akashdeep Singh Virk

**Author notes:** Corresponding author. Sabitoj Singh Virk. Both authors contributed equally to this work.

## Abstract

Formulation and pharmacokinetic liabilities remain major bottlenecks in monoclonal antibody development. Here, we present the agnostic context-embedding transformer (ACeT), an interpretable machine-learning framework that integrates heterogeneous early assay panels into endpoint-specific developability models. Using published antibody datasets, ACeT predicted high-concentration viscosity from four dilute-solution assays with held-out R² ≈ 0.75 and root-mean-square error ≈ 4.8 cP across a 5-45 cP range. From four clearance-related in vitro assays, it predicted mouse intravenous exposure with held-out R² = 0.80 and normalized root-mean-square error = 0.15. On a public 152-antibody panel, ACeT predicted hydrophobic interaction chromatography retention time, an orthogonal stickiness/hydrophobicity readout, with out-of-fold Pearson r² ≈ 0.83 and outperformed a published quantitative structure-property relationship baseline. In an exploratory retrospective analysis using five early developability assays, ACeT classified clinical outcomes (Approved vs Terminated) with balanced accuracy of ∼0.78 on a held-out internal set of 23 clinical IgG1 antibodies with outcome-locked labels and 0.83 on a temporally independent external cohort of 14 antibodies. Feature attribution recovered mechanistically plausible drivers, including the diffusion interaction parameter *k*D and size-exclusion chromatography peak-shape metrics for viscosity and heparin-, baculovirus particle-, and poly-D-lysine- signals for exposure. These results show that routine assay panels can support practical, interpretable machine-learning guided triage of antibodies for formulation and pharmacokinetic risk, and may capture a developability-linked component of downstream progression risk.

## 1. Introduction

Monoclonal antibodies (mAbs) now anchor standard-of-care across oncology, immunology, and neurology, yet their downstream value increasingly depends on whether they can be formulated, delivered, and maintained at exposures compatible with clinical use. Although mAbs generally progress better than small molecules, Phase I-to-approval success remains limited and late-stage failures continue to dominate the fully capitalized cost of development [1–7]. Early triage therefore has to move beyond potency alone and ask a more developability- and delivery-relevant question: which molecules are likely to remain viable once formulation, route-of-administration, and pharmacokinetic constraints are taken seriously?

Among developability liabilities, two are especially consequential for antibodies. First, high solution viscosity at clinically relevant concentrations can render otherwise promising molecules difficult or impractical to formulate for subcutaneous administration. Because subcutaneous delivery is constrained by injection volume, device performance, injectability, and patient tolerance, high-dose antibodies often have to be formulated at concentrations at or above ∼100 mg mL^-1^, precisely the regime where weak protein-protein attractions, reversible self-association, and colloidal crowding can drive steep increases in viscosity [8–15]. Second, unexpectedly rapid clearance reduces systemic exposure and can force impractical dosing schedules or dose escalation. In vitro surrogates such as heparin chromatography, polyspecificity-type assays, and FcRn-related measurements increasingly reveal how nonspecific interactions, charge-mediated liabilities, and recycling behavior shape pharmacokinetics (PK) [16–22]. Together, these risks sit at the interface of developability, formulation, and pharmacokinetics: they determine not only whether an antibody is biophysically tractable, but also whether clinically practical dosing is achievable.

To surface such liabilities before the clinic, discovery groups now deploy multiplex developability panels spanning self-association, hydrophobicity, charge, stability, nonspecificity, and aggregation [23–29]. The rationale is clear: liabilities rarely cluster along a single axis, and no individual assay adequately captures the balance among manufacturability, formulation robustness, delivery feasibility, and PK risk. Recent delivery-focused work has framed this problem in precisely those terms, emphasizing the physicochemical and physiological properties of the subcutaneous injection site, the fate of biotherapeutics after administration, and the need for predictive preclinical tools for absorption and bioavailability [30–35]. Single-assay thresholds remain useful as conservative red flags, but their performance is highly context dependent, and fixed cutoffs rarely provide the sensitivity-specificity balance needed for confident go/kill decisions across diverse molecules and assay panels [22,23,28,29]. What is increasingly needed are integrative frameworks that can learn from semi-orthogonal assay batteries without discarding the experimental context that makes those data informative.

This is where current computational approaches remain limited. Sequence- and structure-based antibody models have advanced affinity optimization, representation learning, and in silico property prediction [36–42], but they generally do not ingest assay context, buffer composition, formulation variables, or experimentally measured readouts that often govern high-concentration behavior, nonspecific binding, clearance, and route-of-administration feasibility. At the same time, public developability datasets remain intrinsically scarce. Assay-rich antibody datasets are expensive to generate, often unpublished, and difficult to standardize across organizations, while sequence disclosures alone rarely preserve the formulation or screening context needed for developability modeling [43–46]. Methods for this space therefore have to be data-efficient, assay-aware, and robust in low-n settings rather than scaled in the manner typical of mainstream machine learning.

Here we introduce the agnostic context-embedding transformer (ACeT), an attention-based framework designed to fuse heterogeneous early-stage assay readouts into a shared latent representation and map that representation to endpoint-specific predictions. In the present study, we apply ACeT to four antibody-relevant settings using published datasets: prediction of high-concentration viscosity from dilute-solution assays; prediction of mouse intravenous exposure from clearance-related in vitro assays; prediction of hydrophobic interaction chromatography retention time as an orthogonal stickiness/hydrophobicity readout; and an exploratory retrospective, developability-linked classification of clinical outcomes from early assay profiles. ACeT is built to reduce experimental burden in early developability by converting small, routinely generated assay panels into quantitative predictions of formulation and pharmacokinetic risk, so that resource-intensive measurements (e.g., high-concentration formulation work and in vivo PK) can be reserved for a smaller, higher-quality subset of candidates. The central question is whether routine early-stage assays, when modeled jointly rather than one at a time, can better identify molecules at risk for concentrated-formulation viscosity, unfavorable systemic exposure, hydrophobicity-linked liabilities, or a broader developability-linked component of progression failure.

Our results indicate that they can. Across formulation-relevant, pharmacokinetic, and progression-linked endpoints, ACeT outperforms single-assay heuristics and conventional baselines while retaining assay-level interpretability and operating within the low-n regime typical of antibody developability datasets. Practically, this shifts early panels from being isolated pass/fail filters to acting as integrated predictors of downstream risk, enabling earlier downselection and more targeted follow-up work. In that sense, ACeT is best viewed as an assay-integrated developability decision model: it learns multivariate relationships across semi-orthogonal readouts and translates routine early measurements into actionable estimates of formulation and PK liability that can streamline candidate ranking and reduce unnecessary experimentation.

## 2. Methods

### 2.1. Dataset preparation and train-test splitting

#### 2.1.1. Viscosity dataset preparation

The high-concentration viscosity corpus comprised 82 IgG1 monoclonal antibodies [47] assayed across 110-200 mg mL⁻¹ (most at ∼150 mg mL⁻¹). Quality control removed seven entries, three statistical outliers exceeding 100 cP (practical syringeability and statistical stability [8,48]) and four profiles with > 20 % missing assay values, yielding 75 complete descriptor-response pairs for subsequent composite-stratified splitting and model development. To reduce dimensionality at n = 75 and emphasize a practically deployable early-screen panel, we fixed the model input to four dilute-solution assays selected for their strongest univariate association [49–50] with viscosity (absolute Spearman correlation) in the curated corpus: DLS diffusion interaction parameter kD (mL/g), SE-UHPLC main-peak plates (EP), SE-UHPLC main-peak FWHM (min), and AC-SINS spectral shift Δλmax (nm) (Supplementary Table S1). This four-assay input schema was held constant throughout subsequent model development. At inference, only these four measurements are required, while other assays reported in the source work were excluded to limit over-parameterization and emphasize practical deployability.

#### 2.1.2. Mouse-clearance dataset preparation

The clearance corpus comprised 55 IgG1 antibodies extracted from a 2023 murine pharmacokinetic study [51]. AUC_0-672h_ values (ng·h mL⁻¹) were divided by 10^6^ (reporting AUCt in units of 10^6^ ng·h mL⁻¹) to keep numeric scales consistent across tasks; two extreme AUCt outliers (13.4 and 1.57, corresponding to 1.34×10⁷ and 1.57×10⁶ ng·h mL⁻¹) were eliminated, leaving 53 profiles. The four in vitro assays exhibiting the highest absolute Spearman correlation with AUCt were retained [49–50] (Heparin_RT, Heparin_pB_buffer, BVP_high, poly_D_lysine; Supplementary Table S1), after which all features were rank-normalized to N(0,1). Values lying beyond ±5 standard deviations were multiply imputed through a ten-iteration MICE protocol. The target variable (AUCt) was not imputed and was excluded from the imputation model; it was carried through unchanged. All subsequent scaling, synthetic augmentation, and resampling steps were fit strictly within training data.

#### 2.1.3. Composite-stratified algorithm for viscosity and mouse clearance

To derive representative yet non-overlapping partitions for the two regression endpoints, we used a composite-stratified splitting procedure that balances coverage in both assay space and response space. Briefly, the assay matrix was standardized, reduced by principal-component analysis (up to five components), and clustered with k-means (3 ≤ k ≤ 8). Independently, the continuous target (neat viscosity or plasma AUCt) was binned into equiprobable quantile bins (1 ≤ b ≤ 5). Concatenating the cluster label with the target-bin label yields composite strata encoding joint membership in feature and response space; a stratified shuffle split on these strata then produces the final train and test subsets. To avoid degenerate splits in these small datasets, we screened a grid of (k,b) settings and selected a parameterization that minimized (i) the fraction of test samples lacking a near neighbor in the training set (an out-of-distribution proxy) and (ii) the mean train-to-test nearest-neighbor distance. The configuration with the lexicographically minimal (OOD-rate, distance) was retained, resulting in a 70: 30 split for viscosity and an 80: 20 split for clearance and giving markedly lower variance in R^2^ and Spearman ρ than conventional continuous stratification.

#### 2.1.4. Clinical-outcome cohorts

From the 137-antibody biophysical landscape [23,29], we defined an internal (outcome-locked development) cohort and an external (temporally independent follow-up) cohort. The internal (outcome-locked 2022) cohort comprised 112 IgG1 antibodies with a resolved program status (Approved or Terminated/discontinued) on or before October 2022; a single 80:20 train_test_split with stratify=y was performed, and the test partition served as the internal development hold-out. The external cohort comprised the remaining 25 antibodies that were still in Phase I-III at the October 2022 freeze [23,29] and were not used during model fitting. By May 2025, 14/25 had reached a terminal outcome (2 Approved; 12 Terminated/discontinued) and were evaluated intact as an out-of-time set, while 11 remain under observation. For modeling, we retained the five developability assays with the highest absolute Spearman correlation with outcome in the training split: accelerated stability under stress (AS; SEC slope), polyspecificity reagent binding (PSR), AC-SINS, an ELISA-based nonspecific-binding readout, and baculovirus particle binding (BVP) (Supplementary Table S1). A binary classifier was then trained to distinguish Approved from Terminated/discontinued programs (negative class: Terminated/discontinued) and applied unchanged to both the internal hold-out set and the external out-of-time evaluation set.

#### 2.1.5. HIC developability dataset preparation

The HIC retention-time corpus comprised 152 antibodies from a public developability panel [46] reporting HIC retention time (hic_rt_min; non-eluting antibodies assigned the 50-min ceiling) together with an orthogonal assay battery and four structure-derived surface patch descriptors (Supplementary Table S1). Because the source study reported a patch-descriptor QSPR fit using the full dataset, we did not define a single train/test split; instead, ACeT was evaluated using 10× repeated 5-fold cross-validation and bagged out-of-fold (OOF) predictions were generated for every antibody (per-antibody OOF predictions averaged across the 10 CV replicates). Three input configurations were tested (assays-only, patch-only, assays + patch); all preprocessing, synthetic augmentation, and any imbalance remediation were confined strictly to the training partition of each fold to prevent leakage. Performance was summarized using continuous regression metrics on bagged OOF predictions (Pearson r², Spearman ρ, RMSE, MAE) and a binary triage analysis at the 30-min cutoff used in the original work (balanced accuracy, MCC; Supplementary Tables S2-S3).

### 2.2. Synthetic augmentation

Synthetic augmentation for the two regression endpoints was performed with the Gaussian Copula Synthesizer (SDV 1.17.4), instantiated with default distribution ‘norm’ so that each empirical marginal was first rank-mapped onto N(0,1) before maximum-likelihood estimation of the correlation matrix; higher-order regularization was disabled to preserve tail dependence among assays. For the viscosity corpus the synthesizer drew one artificial observation per two real samples (1:2), whereas a ratio of 2:1 was adopted for the clearance corpus. These synthetic rows were concatenated with the native training frames prior to any scaling or imbalance remediation, thereby allowing the subsequent scalers to learn from the augmented marginal distributions and eliminating the possibility of leakage from test data. No synthetic augmentation was applied to the clinical-outcome classifier.

### 2.3. Feature transformation, imbalance control and model-selection workflow

After composite-stratified splits had been fixed, the regression and classification pipelines shared a uniform, post-split processing strategy designed to respect the minute sample sizes yet minimize information leakage.

#### 2.3.1. Regression pipelines (viscosity and mouse clearance)

All numerical predictors, native plus Gaussian-copula synthetics, were cast to 32-bit floats and transformed with a Quantile Transformer (1000 quantiles, uniform output) fitted solely on the augmented training matrix; the fitted map was applied verbatim to the held-out test set, ensuring strictly out-of-sample scaling. Within each of the five cross-validation folds the so-scaled training subset was passed to the Imbalanced-Learning-Regression (IBLR 0.2) routine, which automatically chooses between an Edited-Nearest-Neighbours (ENN) under-sampler and a SMOTE over-sampler, or their cascade in either order, according to fold-specific neighborhood densities. This order-agnostic design guarantees that the minority tail of the continuous endpoint is adequately represented while avoiding artefactual duplication when local sparsity is high. Target vectors remained on their original physical scale; a per-fold Quantile Transformer was used only to stabilize the loss surface during training and was inverted once after prediction.

#### 2.3.2. Clinical classification pipeline

For the 112-antibody internal cohort, continuous assay read-outs were variance-stabilized with a Yeo-Johnson Power Transformer fitted on the training partition; no synthetic rows were introduced. Each fold then executed an order-free ENN⇄SMOTE block identical in spirit to the IBLR procedure: ENN first attempted to excise borderline points, the class distribution was re-counted, and SMOTE was applied only if the cleaned minority class still contained at least two examples. The SMOTE k-neighbor parameter was adaptively set to min (5, n_min_−1) to respect the updated class count. Detailed class tallies before and after each operation were printed to the console and logged for audit. Yeo-Johnson PowerTransformer was reused for both the internal test set and the 14-antibody external validation cohort; categorical outcome labels were integer-encoded before one-hot expansion for soft-max training.

#### 2.3.3. Prediction-head benchmarking and ensemble derivation

Five modular heads, multilayer perceptron (MLP), radial-basis function (RBF), Kolmogorov-Arnold network (KAN), cubic spline and pairwise interaction, were benchmarked under an identical five-fold outer cross-validation loop. The head maximizing mean R^2^ (regression) or balanced accuracy (classification) was declared the best head. The five best-head checkpoints (one per fold) constituted an ensemble predictor; test-set estimates were obtained by arithmetic averaging of scalar outputs (regression) or class-probability vectors (classification). This ensembling preserves the variance-reduction benefits of cross-validation while retaining a single, reproducible hyper-parameter configuration for all reported metrics.

### 2.4. ACeT training and evaluation

Each task-specific ACeT model was trained under five-fold cross-validation with shuffled folds. Architectural constants were fixed: 16-dimensional token embeddings, two self-attention heads, a 32-unit feed-forward sub-layer, one transformer block, 0.3 dropout, L_2_=10^−3^ weight decay, and a 64-unit hidden layer in the read-out multilayer perceptron. Optimization employed Adam (initial learning rate 1×10^−3^, mini-batch 32) for ≤1000 epochs; early stopping halted training after 20 stagnant epochs, while ReduceLROnPlateau halved the rate after ten stagnant epochs, with a floor of 1×10^−6^. Regression heads minimized log-cosh loss; the clinical classifier minimized binary cross-entropy. Python, NumPy and TensorFlow seeds were aligned to guarantee bit-wise determinism. In each fold, training used early stopping on validation loss (val_loss) with restoration of the best weights; the five resulting fold-specific models were then averaged to generate the ensemble predictor, which was evaluated once on the held-out test set. Internal hyper-parameters specific to the benchmarked heads (e.g. RBF length-scale) were retained at the empirically optimal defaults established during cross-validation; a future release can substitute this manual tuning with Bayesian optimization via Optuna.

### 2.5. Statistical analysis

Hold-out performance for viscosity and clearance was quantified with R^2^, root-mean-square error, mean absolute error and Spearman’s ρ. Pairwise statistical comparisons for figure-level analyses were performed using the tests indicated in the corresponding figure captions, with multiple-comparison correction where applicable.

### 2.6. Model interpretability

Feature relevance for the viscosity model was quantified with permutation-feature importance: the five-checkpoint ACeT ensemble, exposed through a scikit-learn wrapper that averages predictions and inversely scales them, was evaluated ten times while permuting each input column, and the mean drop in R^2^ served as the importance score. For the mouse-clearance model, KernelSHAP was applied to every checkpoint using 100 representative background points; the resulting tensors (fold × sample × feature) were stacked and averaged, and features were ranked by the grand mean of the absolute Shapley values. All raw importance vectors, SHAP matrices and accompanying figures were archived to enable independent verification.

### 2.7. Software and hardware

All code was executed on a Windows 11 Pro (23H2) workstation equipped with a single NVIDIA A100 GPU (40 GB), CUDA 12.1 and cuDNN 8.9. The Python stack comprised TensorFlow 2.15.0 with Keras 2.15.0, tf-kan 0.1.0, scikit-learn 1.4.1 post1, imbalanced-learn 0.12.2, SDV 1.17.4 (Copulas 0.12.1, RDT 1.14.0, SDMetrics 0.18.0) and SHAP 0.45.0. Core numerics relied on NumPy 1.23.5, SciPy 1.15.2 and pandas 2.2.2, while visualizations were drafted in Matplotlib 3.9.2 and finalized in MATLAB R2022b.

## 3. Results

### 3.1. Overview of ACeT for integrated assay-based developability prediction

We designed the agnostic context-embedding transformer (ACeT) to integrate heterogeneous early-stage assay readouts into a shared latent representation for antibody developability prediction (Fig. 1a-c). Each raw assay value x_i,_ whether continuous, categorical, binary or any other data type, is first projected into a low-dimensional value embedding and combined with a learned feature-identity embedding, yielding one token per measurement (Fig. 1a). The resulting token set is then processed by a compact pre-norm transformer encoder with multi-head self-attention and nonlinear feed-forward blocks, and global average pooling generates a latent representation for downstream prediction (Fig. 1b). In practice, this pooling step is permutation-insensitive once features are aligned to their learned identities; accordingly, all datasets were supplied in the canonical feature order used during training and evaluation.

**Figure 1:**
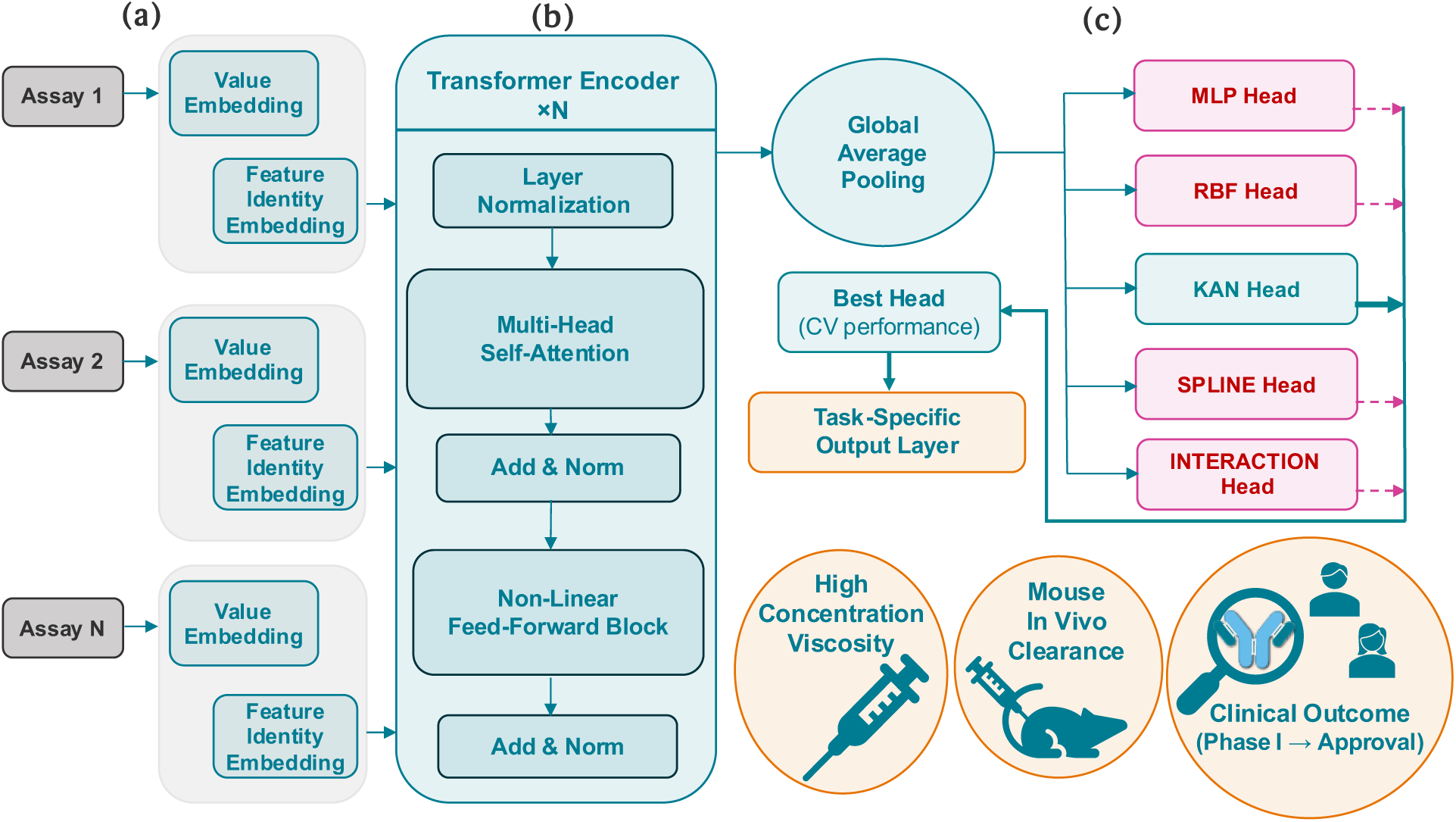
Architecture of the agnostic context-embedding transformer (ACeT) for integrated assay-based developability prediction. **a,** Each assay readout is encoded as the sum of a value embedding and a learned feature-identity embedding, generating one token per measurement. **b,** A compact stack of pre-norm self-attention layers followed by global average pooling converts the token set into a shared latent representation. **c,** Candidate prediction heads with different inductive biases are benchmarked on identical data splits, and the best-performing head is routed to the endpoint-specific output layer, enabling adaptation to viscosity, mouse exposure, HIC, and clinical progression tasks. **Figure 1 Alt Text: Diagram of the ACeT workflow for antibody developability prediction.** The left panel shows tokenization of assay measurements using value embeddings and learned feature-identity embeddings. The center panel shows a compact transformer encoder with self-attention and global average pooling that produces a shared latent representation. The right panel shows task-specific output heads used for regression and classification across multiple developability endpoints.

At the encoder level, ACeT is designed to be flexible with respect to assay composition. It can ingest heterogeneous numeric measurements from experimental or computational pipelines without manual redesign of the core architecture, making it well suited to developability settings in which assay panels evolve across studies or applications. In the present work, the inputs consisted of compact tabular assay panels, and the encoder was intentionally kept lightweight because antibody developability datasets are sparse and a substantially larger model would add capacity without clear benefit while increasing overfitting risk. Although the same tokenization strategy could in principle accommodate additional numeric descriptors or externally derived embeddings when warranted, the focus here is deliberately on routine assay readouts that are practical for early-stage developability screening.

Downstream of the shared encoder, ACeT uses a modular prediction stage in which multiple candidate heads, each expressing a different inductive bias, operate on the same latent representation (Fig. 1c). Because these heads attach only at the representation level, they can be added, removed, or replaced without altering the encoder itself. A lightweight evaluation routine benchmarks candidate heads on identical cross-validation splits and routes the best-performing option to the endpoint-specific output layer. This design allows the same core architecture to support both regression and classification tasks while ensuring that comparisons between heads reflect inductive bias rather than differences in data exposure. Across this study, the resulting framework was applied to high-concentration viscosity prediction, mouse intravenous exposure, HIC retention time prediction, and clinical progression classification.

Taken together, these design choices: feature-specific tokenization, a compact attention-based encoder, pooling to a shared latent representation, and task-tailored prediction heads, are intended to provide useful inductive bias in a setting defined by short assay token sequences and limited sample sizes [52]. ACeT therefore serves as the common modeling framework for all subsequent analyses, enabling assay-level integration across formulation-relevant, colloidal, pharmacokinetic, and progression-linked developability endpoints.

### 3.2. Prediction of high-concentration viscosity from dilute-solution biophysical assays

Using ACeT, we predicted high-concentration solution viscosity from a minimalist panel of four low-concentration developability readouts curated from published measurements [47]: (i) the DLS diffusion interaction parameter kD (mL/g), (ii) SE-UHPLC monomer peak plate count (EP), (iii) SE-UHPLC monomer peak full-width at half-maximum (FWHM, min), and (iv) AC-SINS Δλmax (nm) (Supplementary Table S1). These four variables constituted the full model input, whereas the output was neat viscosity (cP) measured at 110-200 mg mL⁻¹ (most near ∼150 mg mL⁻¹) [47]. On the held-out set of 23 antibodies, predicted and measured viscosities tracked closely across an approximately 5-45 cP range (Fig. 2a).

**Figure 2:**
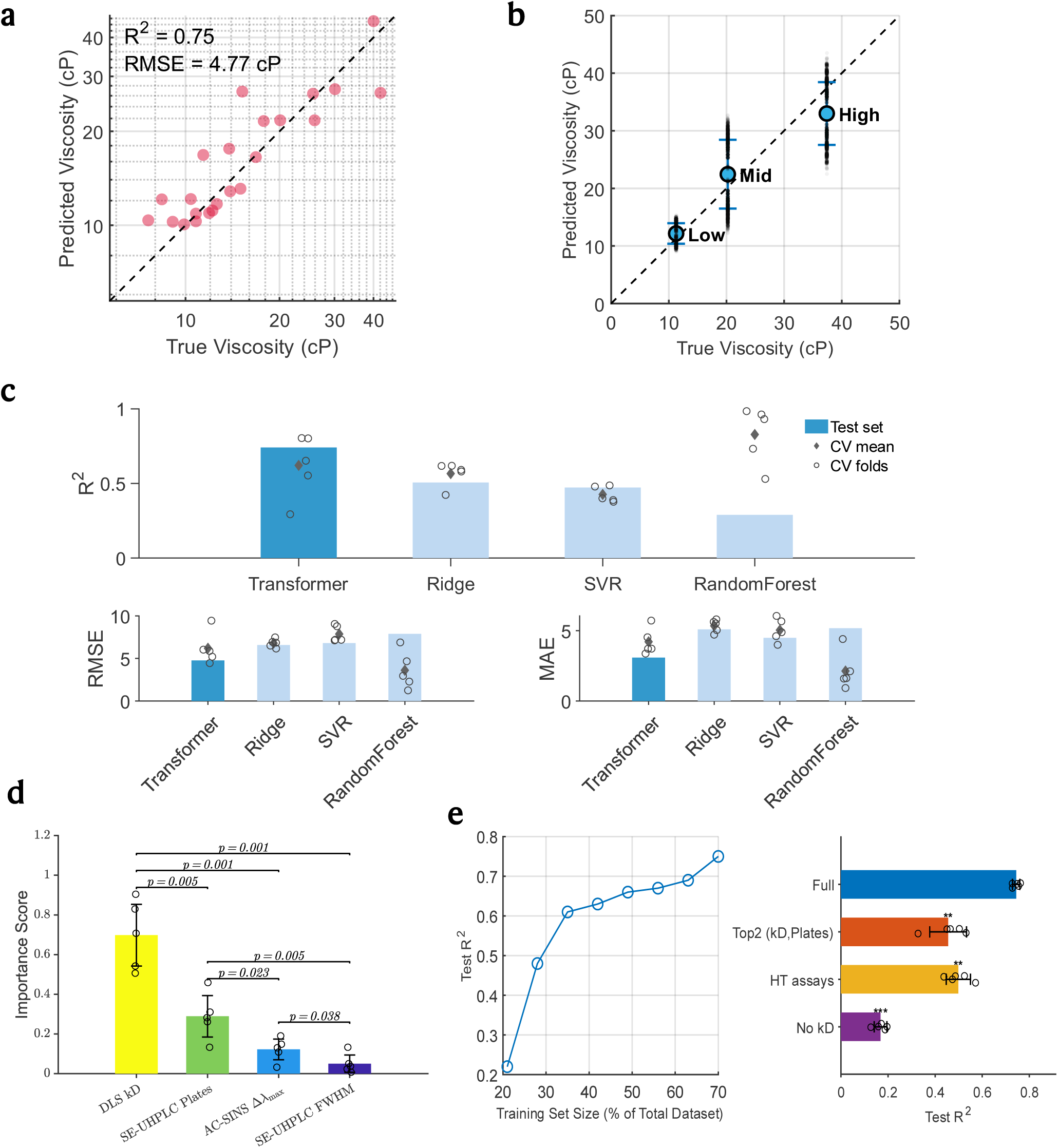
ACeT infers high-concentration viscosity from low-concentration assays. **a,** Parity plot for 23 test antibodies (unity line shown; R^2^=0.75, RMSE ≈ 4.8 cP). **b,** Calibration by viscosity tertile: circles, mean true vs. predicted; vertical bars, bootstrap RMSE (1000 draws); faint dots, resamples; dashed line, unity. **c,** Model comparison (transformer, ridge, SVR, random forest): bars, test metrics; diamonds, five-fold CV means; open circles, CV folds. **d,** Permutation importance of the top four assays: bars, means ± s.d.; open circles, fold values; brackets, pairwise P values (Holm-Šidák). **e,** Left, learning curve of test R^2^ versus training-set size. Right, feature-ablation impact on test R^2^: horizontal bars, mean ± s.d. across five runs; open circles, individual runs; asterisks, significance versus full model (* < 0.05, ** < 0.01, *** < 0.001; paired two-tailed *t*-test, Holm-corrected). **Figure 2 Alt Text: Multi-panel summary of viscosity-prediction performance.** Predicted and measured viscosities from roughly five to forty-five centipoise show close agreement, with R² near 0.75 and RMSE about 4.8 cP. Calibration across low-, mid-, and high-viscosity ranges reveals no systematic bias. The transformer model outperforms ridge, support-vector, and random-forest regressors in both test and cross-validation results. Feature-importance analysis identifies the dynamic-light-scattering interaction parameter kD as the most influential variable, followed by the SEC plate-count metric, while other assays contribute modestly. Model accuracy improves steadily with additional training data, and removal of kD causes a pronounced drop in R². Together, the results highlight ACeT’s predictive strength and the main assay drivers of viscosity.

Most points fell near the unity line, yielding an overall coefficient of determination R² ≈ 0.75 and a root-mean-square error ∼4.8 cP on the test set. The model maintained strong performance even at the extremes, showing minimal error in the low-viscosity (∼5-15 cP) and mid-viscosity (∼15-30 cP) ranges, and only a slight underestimation bias for the highest-viscosity samples (>30 cP), all within experimental error. Because the source measurements were collected at varying protein concentrations between 110 and 200 mg mL^-1^ without explicit normalization [47], we interpret the model within this concentration window rather than as a concentration-adjusted viscosity predictor. Even under that constraint, the held-out performance indicates that dilute-solution assay features retain substantial information about concentrated-solution behavior.

Stratifying predictions by viscosity regime confirmed this robustness (Fig. 2b). In the low, medium and high bins, the transformer’s estimates remained tightly clustered around the unity line, and the surrounding bootstrap cloud showed no systematic deviation, indicating that the model does not rely on interpolation within any single range. By capturing nonlinear assay-viscosity relationships without systematic deviation across the viscosity spectrum, the transformer outperforms conventional approaches.

ACeT also outperformed baseline machine-learning models in both cross-validation and independent tests (Fig. 2c). For example, on the test set the transformer achieved R² >0.7, substantially higher than the best alternative (ridge regression, R² ∼0.5) and well above support-vector regression (∼0.47) or a random forest (∼0.29). This translated to markedly lower prediction errors: the transformer’s test RMSE was ∼5 cP, versus ∼7-8 cP for the baselines, and its mean absolute error (∼3 cP) was about 2 cP lower than those of conventional models.

Notably, the transformer’s performance generalized consistently (mean R² ∼0.62 across folds, nearly matching the independent test R²) with no signs of overfitting, whereas the random forest exhibited high variance and severe overfitting (some cross-validation folds showed spuriously high R² up to ∼0.9, but overall test R² dropped to ∼0.3; Fig. 2c). These results highlight the ACeT’s ability to learn generalizable viscosity predictors from limited data, unlike simpler linear models that under-fit or complex ensembles that overfit. We included ridge, SVR and random-forest baselines as domain-standard comparators spanning linear, kernel and tree-ensemble families that remain competitive in low-feature biotherapeutics developability modeling [53–55].

Permutation analysis indicated that the model relied on physically interpretable assay features. The dynamic light scattering interaction parameter kD emerged as the dominant contributor (Fig. 2d), with a substantially larger importance score (∼0.7 on a 0-1 scale) than any other input. This parameter, measured in dilute solution, reflects net protein-protein interaction propensity: more negative kD values indicate stronger attractive interactions that often portend high viscosity [13,56]. Consistent with domain knowledge, antibodies with unfavorable (negative or low) kD indeed showed elevated viscosity [13,56,57], and the model appropriately up-weighted this feature.

Notably, the second most important predictors were two SEC/SE-UHPLC monomer peak-shape descriptors: plate count (peak sharpness/efficiency) and FWHM (peak width) (Fig. 2d). In chromatography, plate count and FWHM quantify band broadening (higher plates and lower FWHM correspond to narrower peaks) [58]. For proteins, apparent peak broadening and asymmetry can reflect sample heterogeneity (e.g., low levels of higher-order species) but can also arise from reversible self-association during the separation or from secondary (non-size) interactions with the stationary phase [59,60]. Because the same weak, transient intermolecular attractions that promote reversible self-association at dilute concentration are also implicated in cluster formation and steep viscosity increases at >100 mg mL⁻¹ [10,13,57], the observed SEC-viscosity coupling is biophysically plausible. A high plate count (sharp monomer peak) correlated with lower viscosity [47], and accordingly this feature received a moderate importance score (∼0.3), while AC-SINS Δλmax contributed additional orthogonal information [61]. In the dataset [47], plate count and FWHM are inversely related measures of peak dispersion and were highly correlated; accordingly, the model relied most strongly on plate count, rendering FWHM partly redundant under permutation. Overall, the learned dominance of kD and SEC peak-shape metrics aligns with established protein-interaction drivers of high-concentration viscosity, reinforcing interpretability and credibility.

Finally, learning-curve and ablation studies underscored data-efficiency and the critical nature of kD (Fig. 2e). Model performance improved monotonically with training set size and showed no sign of plateau within the available data, indicating that additional training examples would likely further enhance accuracy. In parallel, removing the DLS kD feature, identified above as critical, caused the test-set R² to plummet from ∼0.75 to ∼0.17 (“No kD”), a highly significant loss of predictive power (p < 0.001). Using the two most important features (kD and SEC plate count) only partially recovered performance (R² ≈ 0.45, “Top2”), and relying solely on the high-throughput assays (omitting SEC) improved only slightly (R² ≈ 0.49, “HT assays”). Together, these tests confirm that while all assay inputs contribute, the DLS kD metric is indispensable for capturing the bulk of the viscosity signal. Consistent with prior single-descriptor viscosity screens based on kD [13], a simple kD-only baseline regression captured only modest variance on this dataset (test R² ≈ 0.27), whereas adding SEC peak-shape information improved baseline performance (Supplementary Table S4), and the full ACeT model further increased accuracy by integrating all four assays nonlinearly. Moreover, the steadily climbing learning curve suggests the ACeT model is not data-saturated and would benefit from larger training datasets, implying that its current performance represents a balance limited by available data rather than an intrinsic ceiling.

### 3.3. Prediction of mouse intravenous exposure from in vitro clearance-related assays

Extending the ACeT framework to pharmacokinetics, we trained the architecture to map a focused set of four in vitro PK assay read-outs (Heparin chromatography retention time (Heparin_RT), Heparin %B at elution (Heparin_pB_buffer), Baculovirus particle binding at high concentration (BVP_high), and poly-D-lysine binding) directly into quantitative predictions of mouse exposure, reported as the area under the plasma concentration-time curve from time 0 to the last sampled timepoint (AUC_0-t_, hereafter AUCt) [51]. Here AUCt corresponds to AUC_0-672h_ (as reported in the source study [51]) and is expressed in units of 10^6^ ng·h mL⁻¹ (i.e., AUC_0-672h_ divided by 1,000,000), matching the values reported in Fig. 3 and Supplementary Table S1.

**Figure 3:**
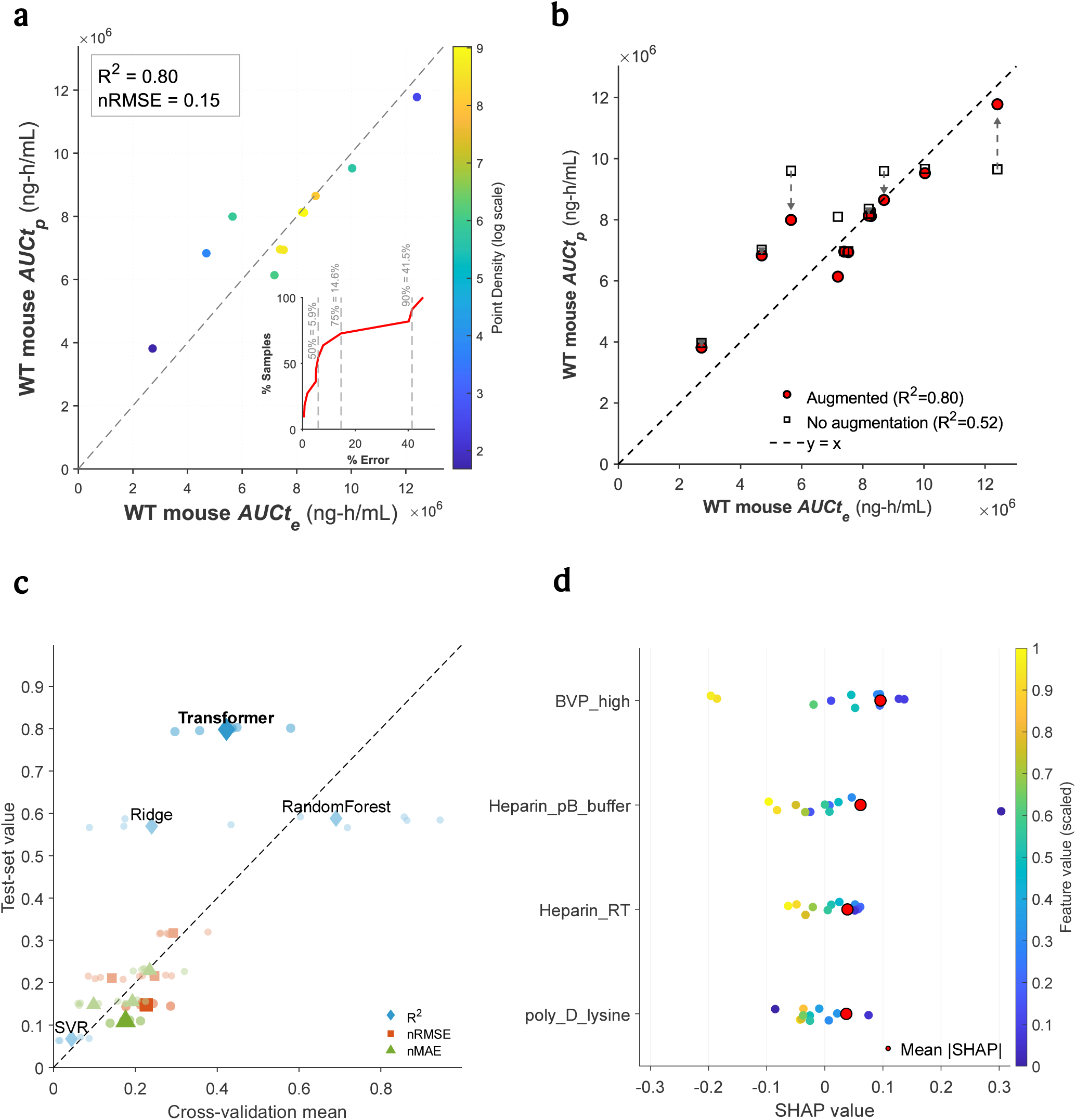
ACeT predicts mouse intravenous exposure from in vitro clearance-related assays. **a,** Density-colored parity plot for the 11-antibody test set (dashed line, unity; R^2^=0.80, nRMSE ≈ 0.15); inset, cumulative distribution of absolute-percentage error. **b,** Effect of synthetic augmentation: red circles, augmented model (R^2^=0.80); open black squares, unaugmented (R^2^=0.52); dashed arrows link paired predictions for each antibody, highlighting the systematic improvement. **c,** Generalization comparison of transformer, ridge, SVR and random-forest regressors. Diamonds, R^2^; squares, nRMSE; triangles, nMAE; each marker plots cross-validation mean (x-axis) against test-set value (y-axis); dashed diagonal, y = x. **d,** KernelSHAP beeswarm for the trained transformer; dots, individual SHAP values colored by scaled feature level; red circles, mean |SHAP|, identifying the four dominant assays (BVP_high, Heparin_pB_buffer, Heparin_RT, poly_D_lysine). **Figure 3 Alt Text: Four panels show mouse intravenous-clearance prediction based on in vitro assays.** Predicted and observed area-under-the-concentration-time-curve values display strong agreement, with R² about 0.80 and normalized RMSE near 0.15; most errors are below fifteen percent. Data augmentation markedly reduces prediction error relative to the unaugmented model. The transformer generalizes better than ridge, support-vector, and random-forest regressors, achieving higher R² and lower error metrics. SHAP analysis highlights BVP_high, heparin chromatography measurements, and poly-D-lysine binding as the major contributors, where stronger signals correspond to faster intravenous clearance. Together, the results highlight both the predictive accuracy and mechanistic interpretability of the clearance model.

Using paired in vitro/in vivo data, ACeT was optimized by cross-validation and then evaluated on a held-out test set. Predicted and observed AUCt values agreed closely across an approximately 20-fold exposure range (Fig. 3a), yielding test-set R² = 0.80 and nRMSE = 0.15. Most antibodies were predicted with high fidelity: the median absolute percentage error was ∼5.9%, and 75% of test compounds fell within 15% of the measured AUCt (Fig. 3a, inset). Only a small minority showed larger deviations (90th percentile error ∼41.5%). Given that the inputs were limited to four in vitro assay readouts, this level of agreement indicates that ACeT captured a meaningful in vitro-to-in vivo relationship for mouse clearance behavior.

Because the original dataset was small, we augmented the training data by fitting a Gaussian-copula model to the training partition only and sampling additional synthetic in vitro assay profiles with realistic marginal distributions and correlation structure, thereby enlarging and regularizing the training distribution. The held-out test set contained only real (unaugmented) antibodies. Without augmentation, ACeT overfit the limited data and generalized poorly (R² ≈ 0.52; black squares in Fig. 3b). In contrast, the model trained with augmentation achieved R² = 0.80 on the same test compounds (red circles in Fig. 3b), with predictions consistently closer to the unity line.

Overall, augmentation reduced error substantially, increasing R² from ∼0.52 to 0.80 and lowering nRMSE from ∼0.25 to 0.15. By broadening the effective training distribution while preserving a strictly real test set, augmentation reduced sensitivity to sampling noise and enabled more stable learning of the in vitro–in vivo relationship in this low-n regime.

ACeT generalized far better than baseline models (Fig. 3c). Although its cross-validation R^2^ was modest (0.42), training on the full dataset and testing on unseen compounds yielded R^2^=0.80 with the lowest errors (nRMSE = 0.15; nMAE = 0.10). The held-out test set covers a comparable AUCt range to the training data (272-1241 vs 182-1318; SD 259 vs 244), so the higher test R^2^ is consistent with training the final ensemble on the full augmented data rather than an unduly easy split. Because each cross-validation model was fit on ∼80% of a small, high-dispersion dataset, the out-of-fold R^2^ represents a conservative estimate; ensembling models trained on the full augmented set recovers additional signal, yielding R^2^ =0.80 on unseen compounds. Importantly, the scale-normalized errors (nRMSE, nMAE) were lowest on the test set, indicating genuine error reduction rather than variance-driven inflation of R^2^. In contrast, a Random Forest regressor overfit (R^2^ =0.58), Ridge regression underfit (R^2^ =0.56), and SVR underperformed in both settings (R^2^ <0.1). Thus, among the models tested, ACeT provided the most reliable mapping from in vitro assay profiles to in vivo exposure.

To clarify ACeT’s decision-making and verify alignment with established pharmacokinetic drivers, we applied SHAP (Shapley additive explanations) [62] to the trained model. High values of BVP_high (Baculovirus Particle, high concentration, binding assay) or Heparin_pB_buffer (Heparin percent B buffer elution assay) consistently yielded negative SHAP contributions (yellow points left of center), indicating that strong signals in these assays drive lower predicted AUCt (i.e. faster clearance). Conversely, low feature values (purple points) shifted SHAP contributions to positive, reflecting higher exposure for compounds with minimal binding or clearance in those assays. Heparin_RT (Heparin chromatography retention time) followed the same trend, strong heparin interactions were associated with reduced AUCt, consistent with accelerated clearance of highly bound molecules. This pattern accords with pharmacological intuition: both assays capture a propensity for nonspecific binding or rapid metabolism, traits that inversely correlate with systemic exposure [22,63]. Finally, poly_D_lysine, an assay for nonspecific binding to a cationic surface, exhibited the same relationship: compounds that bind strongly (high feature value) were predicted to have lower AUCts, whereas weak binders retained higher exposure [47,51].

Overall, the mouse-IV exposure model combined strong held-out accuracy, clear benefit from regularized training, and assay-level interpretability. These features make ACeT a practical in vitro/in vivo prioritization tool for identifying candidates more or less likely to show favorable systemic exposure before resource-intensive in vivo PK studies, with potential value for streamlining early antibody developability and reducing unnecessary animal use in keeping with the 3Rs.

### 3.4. Clinical progression classification from early developability assays and retrospective portfolio analysis

Using five early-stage developability assays curated from historical clinical candidates [23,29]: accelerated stability under stress (AS; SEC slope), polyspecificity reagent binding (PSR), AC-SINS, an ELISA-based nonspecific-binding readout, and baculovirus particle binding (BVP) (Supplementary Table S1), we trained ACeT to classify Phase I antibodies as eventually Approved or Terminated. Cross-validated training did not indicate material overfitting, and the held-out internal test set of 23 antibodies yielded 76.9% sensitivity and 80.0% specificity (10/13 approvals identified; 8/10 terminations correctly flagged), corresponding to a balanced accuracy of ∼78% (Fig. 4a). In practical terms, the model recovered roughly three-quarters of eventual approvals while filtering out four-fifths of eventual failures. The selected operating point, determined by maximizing the Youden index (sensitivity + specificity – 1), lay close to the default probability threshold of 0.5 and produced only three false-negative approvals and two false-positive terminations.

**Figure 4:**
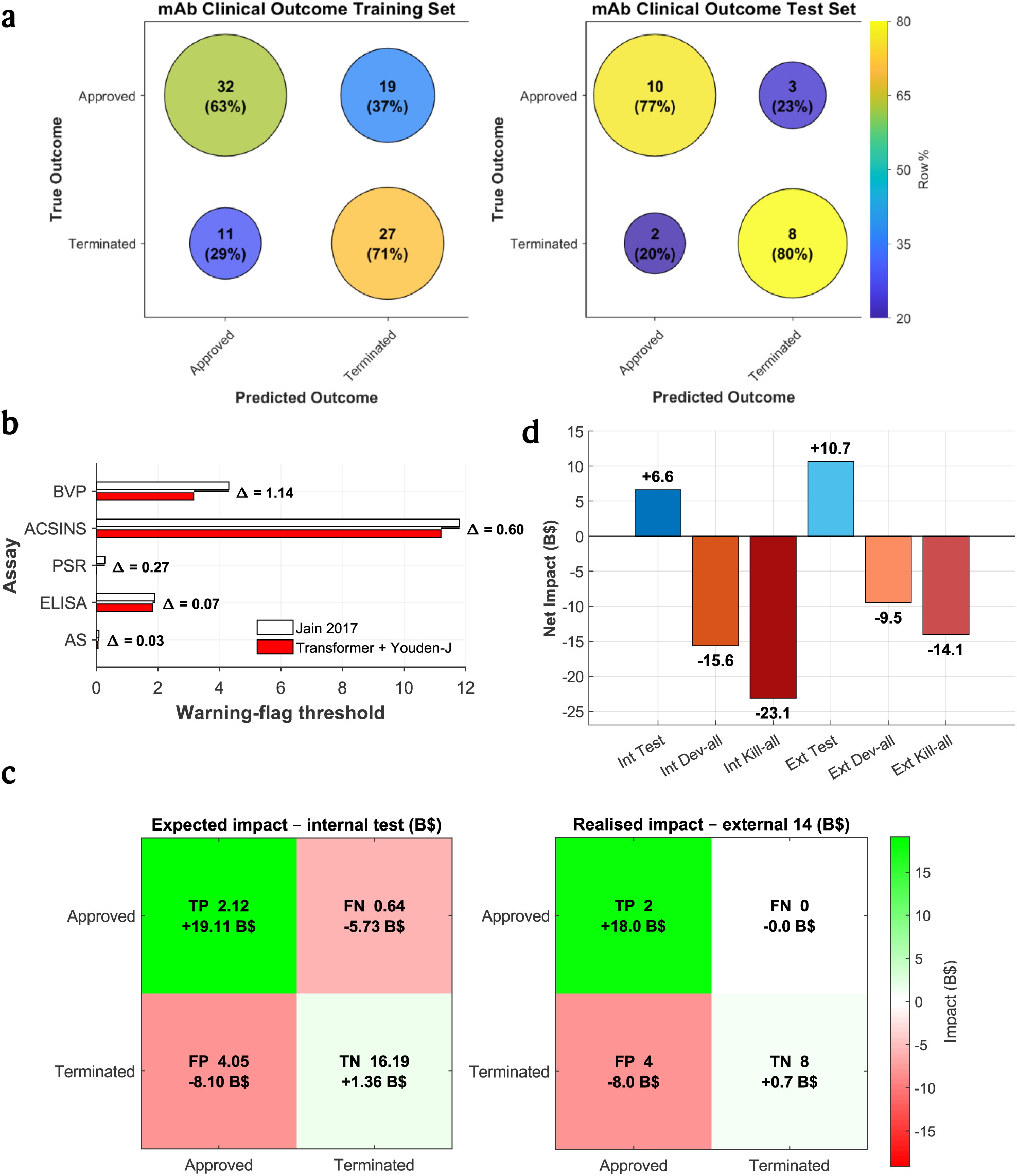
ACeT classifies Phase I antibodies from early developability assays and contextualizes classifier performance with retrospective portfolio analysis. **a,** Bubble confusion matrices: left, training set; right, held-out internal test set (*n* = 23). Bubble area scales with sample count and color with row percentage. **b,** Single-assay ‘red-flag’ thresholds from the 2017 rule set (white) versus ACeT’s Youden-optimized cut-offs (red); horizontal brackets annotate the offset Δ. **c,** Heat maps converting classifier outcomes to illustrative net-present-value consequences under the stated assumptions for the internal portfolio illustration (left) and the 14-mAb external cohort (right). **d,** Portfolio-level NPV for three illustrative policies, AceT triage, develop-all and kill-all, applied to the internal portfolio illustration (left trio) and the external cohort (right trio); bars are labeled with net billions (US$). **Figure 4 Alt Text:** Bubble confusion matrices summarize ACeT classification of Phase I antibodies in training and held-out internal test sets, showing about 77% sensitivity and 80% specificity on the internal test cohort. Comparison with single-assay red-flag thresholds shows that the integrated classifier achieves a more favorable balance of sensitivity and specificity than fixed assay cutoffs. Heat maps and bar charts translate classifier outcomes into an illustrative retrospective portfolio analysis for the internal portfolio illustration and an external 14-antibody cohort under the stated economic assumptions.

This endpoint should be interpreted as a developability-linked classification task rather than a comprehensive estimate of approval probability. Clinical progression depends on efficacy, safety, trial design, indication, and strategic considerations in addition to developability [64]. Nevertheless, early assay profiles can influence downstream success by constraining manufacturability, formulation and delivery feasibility, and achievable exposure through nonspecific interactions and clearance-related liabilities [8–11,16–22,48]. Consistent with that view, prior surveys of the clinical-stage antibody landscape have reported enrichment of developability liabilities among programs that fail to progress [23,29]. ACeT should therefore be viewed as estimating a developability-linked component of progression risk that complements, rather than replaces, efficacy and safety assessment.

We next benchmarked ACeT against literature-derived single-assay rules to place the classifier in context (Fig. 4b). Previous studies [23,29] proposed “red-flag” thresholds for individual biophysical readouts, such as PSR > 0.27 or AC-SINS > 11.8 nm, as indicators of developability risk. Using the published rule set [23] as a conservative one-flag heuristic, any antibody crossing at least one suggested cutoff captured many eventual failures but also flagged a substantial number of antibodies that ultimately progressed. Tightening the criterion by requiring multiple assays to exceed threshold improved specificity only by sacrificing recall. By contrast, ACeT learns a multivariate decision boundary across the full assay panel and, at its selected operating point, achieved ∼77% sensitivity and 80% specificity on the held-out internal test set, a balance that no single-assay rule examined here matched (Fig. 4b). These results indicate that jointly modeling semi-orthogonal developability signals improves early separation between programs more and less likely to progress beyond what is possible with isolated assay cutoffs.

To translate classifier behavior into operational terms, we then carried out a retrospective portfolio simulation using published estimates of phase-transition priors, development costs, and launch value (Table 1). We treat this analysis as contextualization of classifier performance rather than as a primary claim of the study. For the internal portfolio illustration, we applied the observed sensitivity and specificity to a hypothetical 23-program Phase I portfolio using a 12% baseline success prior [1,4]. Under these assumptions, the expected cell counts were TP = 2.12, FN = 0.64, FP = 4.05, and TN = 16.19, with corresponding net-present values of +$9 B for a launch [65], -$9 B for a missed launch [7,65], -$2 B for a continued failure [7], and +$0.084 B for an early kill [7] (Table 1). This yielded the color-scaled heat-map values shown in Fig. 4c (left): +$19.11 B for TP, -$5.73 B for FN, -$8.10 B for FP, and +$1.36 B for TN, corresponding to an expected net of +$6.6 B for ACeT-guided triage. Under the same parameterization, a develop-all policy yielded -$15.6 B and a kill-all policy -$23.1 B (Fig. 4d). Thus, within this retrospective framework, improved control of both false positives and false negatives translated into a favorable portfolio ranking under the stated assumptions.

**Table 1.** Illustrative economic assumptions used for retrospective portfolio analysis of ACeT Phase I classifications. For the internal portfolio illustration, a Phase I probability of success (PoS = 12%) [1,4] was applied to a hypothetical 23-program portfolio and combined with ACeT’s observed sensitivity and specificity to obtain expected cell counts (for example, TP = 23 × 0.12 × 0.769 ≈ 2.12). Each outcome was then assigned a net-present value: +$9 B for a launch (TP) [65], -$9 B for a missed launch (FN) [7,65], -$2 B for a continued failure (FP) [7], and +$0.084 B for an early kill (TN) [7]. These assumptions were used to generate the heat-map analysis in Fig. 4c.

| Outcome | Abbrev. | Assumed phase-I prior | Per-asset value | Rationale |
| --- | --- | --- | --- | --- |
| Launch success | TP (true positive) | 12 %→fractional 2.12 | +\$9 B | \$11 B fully capitalized mean net-present value (NPV) minus \$2 B R&D cost |
| Missed success | FN (false negative) | 0.64 | −\$9 B | Opportunity cost of wrongly killing a future launch |
| Continued failure | FP (false positive) | 4.05 | −\$2 B | Sunk Phase II/III spend with no return |
| Early kill | TN (true negative) | 16.19 | +\$0.084 B | Median Phase II/III savings (\$25 M + \$59 M) |

The external evaluation set comprised 14 of the 25 mAbs that were still progressing through Phase I-III at the October 2022 outcome freeze [23,29] and had reached terminal outcomes by May 2025. Two ultimately gained approval (lebrikizumab and olokizumab) [66,67], whereas 12 were terminated or discontinued. Without retraining, ACeT classified this out-of-time set as 2 true positives, 0 false negatives, 4 false positives, and 8 true negatives, corresponding to a balanced accuracy of 0.83 (Fig. 4c, right). Applying the same illustrative cost framework to these realized counts yielded a net of +$10.7 B, compared with -$9.5 B for develop-all and -$14.1 B for kill-all (Fig. 4d). The classifier’s advantage under the stated assumptions was therefore preserved in a temporally independent set of molecules whose outcomes were unresolved when the original dataset was assembled.

Varying the assumed clinical-success prior from 8% to 20% and scaling launch net present value by ±30% changed the absolute dollar values but did not alter the ranking of ACeT-guided triage relative to the develop-all and kill-all comparators (data not shown). We therefore interpret the portfolio analysis as a retrospective way to contextualize classifier behavior under plausible development economics, while the primary result remains that integrated early assay modeling recovered a balanced and operationally useful separation between antibodies more and less likely to progress.

### 3.5. Hydrophobicity/stickiness proxy prediction: HIC retention time

Hydrophobic interaction chromatography (HIC) retention time provides an orthogonal, widely used proxy for antibody hydrophobicity and “stickiness,” properties linked to nonspecific interactions and developability risk. We therefore applied ACeT to a public 152-antibody developability panel reporting HIC retention time (hic_rt_min) together with a multiplex assay battery and four structure-derived surface patch descriptors [46]. Models were evaluated using 10× repeated 5-fold cross-validation and summarized as bagged out-of-fold (OOF) predictions (Methods; Supplementary Tables S2-S3). HIC retention times span 20.18-50 min, with a substantial fraction of antibodies at the 50-min ceiling (set to 50 min in the source study for non-eluting antibodies).

Using assays-only inputs (excluding patch descriptors), ACeT achieved strong continuous prediction performance (bagged OOF Pearson r² = 0.825; Spearman ρ = 0.881; RMSE = 4.995 min; MAE = 3.256 min; Supplementary Table S2; Supplementary Fig. S6). This assays-only model also outperformed structure-derived patch descriptors, and adding patch descriptors to the full assay panel did not improve beyond assays alone (assays + patch Pearson r² = 0.786). In an apples-to-apples patch-only setting, ACeT exceeded a fold-matched refit of the published linear surface-patch QSPR baseline (Pearson r² = 0.585 vs 0.507; RMSE = 7.451 vs 7.763 min; Supplementary Table S2; Supplementary Fig. S7).

When recast as a binary triage at the 30-min cutoff used in the original work, assays-only ACeT reached balanced accuracy 0.891 and MCC = 0.777 (baseline: 0.777 and 0.550; Supplementary Table S2; Supplementary Fig. S8). Full parity plots, replicate-level variability, and per-antibody OOF prediction tables are provided in Supplementary Figs. S6-S8, Supplementary Tables S2-S3, and Supplementary Data.

## 4. Discussion

ACeT shows that routine antibody developability assays can be integrated within a single attention-based machine learning framework to generate practically useful predictors across formulation-, colloidal-, pharmacokinetic-, and progression-linked endpoints. This matters because publicly available antibody developability datasets are typically low-n and context-rich, and the assay readouts reported across studies rarely conform to a single standardized panel. Existing tabular models such as TabNet [68], FT-Transformer [69] and TabPFN [70] have advanced learning on structured data, but they generally assume a defined feature schema and are less naturally suited to settings where the available readouts differ between datasets or endpoints. ACeT is designed for that setting: it can accept heterogeneous assay inputs without manual architectural redesign and route shared assay representations to endpoint-specific heads according to the prediction task. Antibody language models [40,42,71] provide powerful residue-level sequence representations, but they do not natively fuse orthogonal experimental readouts from routine developability screens. ACeT therefore occupies a distinct methodological niche by learning directly from assay panels while remaining compatible with Shapley additive explanations, so that individual predictions can be decomposed into assay-level contributions. That interpretability is important not only for mechanistic confidence, but also for practical deployment in early-stage developability workflows where screening decisions must be justified in physical rather than purely statistical terms [62].

This is clearest in the viscosity setting. For high-concentration viscosity, ACeT concentrates attention on the dynamic light-scattering interaction parameter kD while also drawing heavily on SEC/SE-UHPLC peak-shape features such as plate count and FWHM. That pattern is physically coherent. Increasingly negative kD values are classically associated with attractive self-interactions that promote reversible clustering and steep viscosity rises in concentrated mAb solutions [13,56,57], while peak broadening or asymmetry in SEC can reflect reversible self-association during separation or secondary interactions with the stationary phase [58–60]. In effect, the model combines orthogonal dilute-solution readouts that plausibly report on the same intermolecular forces that become problematic above 100 mg mL^−1^ [10,57]. This explains why ACeT improves substantially over kD-only baselines while retaining mechanistic interpretability (Supplementary Table S4). Rather than replacing established descriptors, it integrates them with additional assay information to yield a more stable predictor of formulation liability. From a developability and delivery standpoint, this endpoint is especially consequential because high viscosity is one of the main barriers to highly concentrated antibody products, particularly for subcutaneous administration where injectability and dose-volume constraints create a practical soft ceiling in the approximately 15-20 cP range [11,30–35,72]. On a contemporary 75-mAb panel, ACeT achieves holdout performance in the mid-0.7 R^2^ range on a test cohort representing roughly one-third of the data, outperforming ridge, random-forest, and support-vector baselines by more than 25% while retaining usable accuracy at both viscosity extremes. In that sense, the model advances beyond earlier machine-learning studies that were trained on only a few dozen antibodies and often reported categorical pass-fail outputs without absolute error metrics [13,73]. The collapse in performance after kD ablation further supports the view that short-range intermolecular interactions remain a dominant determinant of viscosity in the >100 mg mL^−1^ regime [74], while the still-rising learning curve suggests that additional gains are likely if assay-rich viscosity data can eventually be pooled across organizations rather than remaining siloed.

A complementary result emerges for hydrophobicity-related developability risk. Using hydrophobic interaction chromatography retention time as a proxy for stickiness, ACeT achieved strong continuous prediction on the public 152-antibody panel [46] under repeated out-of-fold evaluation and substantially exceeded the published QSPR baseline based on structure-derived hydrophobic patch descriptors. Notably, assays-only ACeT performed as well as or better than models that included patch descriptors, and a patch-only ACeT variant also outperformed the original baseline. This indicates that the gain is not simply due to privileged inputs, but to modeling capacity: the attention framework is better able to capture nonlinear structure in the available data. Together with the kD-dominated viscosity attributions, these findings suggest that routine assay batteries encode multiple, semi-orthogonal colloidal interaction modes, including electrostatically driven self-association and hydrophobic stickiness, and that these modes can be learned jointly within a single assay-aware encoder. That is important because early developability decisions are rarely made on one liability alone. Molecules that are acceptable on one assay can still fail because of the combined effect of several moderate liabilities, and a model that integrates these signals is inherently more useful than a series of disconnected thresholds applied one readout at a time.

In the pharmacokinetic setting, ACeT similarly extracts meaningful signal from a compact in vitro assay panel. When trained on four standard assays, the model provides useful estimates of mouse intravenous exposure and clearance, complementing physiologically based PK approaches that often require much more extensive antibody-specific parameterization [51]. The assay attributions are again mechanistically credible: heparin, BVP, and poly-D-lysine emerge as major contributors, consistent with longstanding evidence that nonspecific and charge-mediated interactions are important drivers of rapid clearance among antibodies that are otherwise similar in canonical developability metrics [22,51,63,75,76]. The model therefore does not eliminate the need for in vivo PK work, but it can help prioritize molecules before animal studies and improve the efficiency with which those studies are deployed. In that sense, the framework is aligned with the practical goal of reducing avoidable animal use while enriching for candidates with a lower probability of early PK failure. The marked benefit of noise-based data augmentation is also notable. In this low-n regime, synthetic perturbation substantially reduces prediction error, echoing broader reports that transformer-style models can benefit from carefully chosen regularization when datasets are too small to support conventional deep-learning scaling [70]. Looking ahead, larger multi-sponsor datasets would likely smooth the remaining heteroscedasticity and improve rank-order stability, and privacy-preserving collaborative schemes such as federated learning and related distributed-learning frameworks offer one plausible route toward that goal [77–79].

The clinical-outcome task should be interpreted in the same pragmatic spirit. The point is not that developability assays determine approval, and the work does not make that claim. Clinical progression also depends on efficacy, safety, indication, trial design, competitive landscape, regulatory strategy, and sponsor-level decisions that lie outside any developability model. What ACeT tests is a narrower and more defensible question: whether an experimentally tractable early assay panel contains a developability-linked signal that enriches for downstream progression among Phase I antibodies. The answer appears to be yes. On the historical set, the model reaches balanced accuracy in the high-seventies and outperforms simple single-threshold heuristics such as fixed cutoffs on polyspecificity or self-interaction assays, which often either eliminate molecules that might have succeeded or allow liabilities to advance unchecked [23,29]. Just as importantly, the out-of-time evaluation on antibodies whose outcomes resolved only in 2025 suggests that the learned decision boundary is not purely retrospective overfitting, but captures a signal that remains relevant as the development landscape evolves. That does not make the endpoint noise-free; far from it. It means only that assay-derived developability information appears to contribute a stable, nontrivial component to downstream progression risk. Framed this way, the result is not an overclaim about approval forecasting, but a strong argument that integrated assay modeling can improve early candidate ranking beyond what is possible with isolated assay cutoffs.

The retrospective economic analysis is best understood as a translation layer. By mapping the classifier’s confusion matrix onto plausible development-cost and asset-value assumptions [6,80], the portfolio simulation shows how improved control of false positives and false negatives could propagate into operational consequences in biologics R&D. The absolute dollar estimates are necessarily assumption-dependent and should therefore be treated as illustrative. The more durable conclusion is that better early selection precision can materially affect how capital and experimental effort are distributed across antibody portfolios when late-stage failures are costly.

More broadly, ACeT argues against a developability strategy built around isolated assay thresholds and static rules. Traditional early-stage triage often treats each assay separately, with fixed decision limits applied in parallel and little attempt to learn how moderate liabilities combine. ACeT replaces that fragmented logic with a learned, interpretable decision surface that integrates heterogeneous biophysical measurements into a single risk-aware framework. In practical terms, this can make early screening more quantitative, more transparent, and potentially more efficient. A molecule that looks borderline on several assays may be appropriately deprioritized even if it does not violate any one hard cutoff, while another that fails a single heuristic threshold may be retained if the broader assay context is favorable. That is a more realistic representation of how developability risk is actually distributed across antibody candidates.

Several considerations frame the present scope. Antibody developability datasets are inherently scarce and experimentally heterogeneous, so models in this domain must be data-efficient and are likely to benefit further from broader assay-rich data sharing. The present study is centered on IgG-like molecules and assays generated under defined protocols, meaning that extension to bispecifics, Fc-engineered formats, new laboratory settings, or emerging assay implementations may benefit from modest recalibration or fine-tuning. In addition, the current work emphasizes early developability, formulation, and PK-related readouts rather than the full manufacturing and delivery continuum; future incorporation of process-level endpoints such as concentrate-ability or UF/DF performance [81] and broader translational datasets should further strengthen the framework.

Despite these considerations, ACeT shows that early-stage assay data can be leveraged more effectively than has generally been demonstrated to date in antibody developability [13,23]. Rather than treating routine biophysical assays as isolated filters, the framework learns how they interact across formulation, colloidal, pharmacokinetic, and downstream progression-relevant dimensions. The result is a practical machine learning approach that shortens the distance between assay generation and actionable ranking, offers mechanistic interpretability at the level of individual readouts, and provides direct delivery relevance by identifying liabilities linked to concentrated formulation behavior, subcutaneous administration feasibility, and clearance risk. In that sense, ACeT is not simply another predictive model for one endpoint. It is a proof of concept that heterogeneous assay panels can support a unified, interpretable, data-efficient developability framework for earlier selection of antibody candidates with improved prospects for manufacturability, delivery, and clinical progression.

## Data and code availability

All raw datasets analyzed in this study were obtained from published sources [23,29,46,47,51]. Supplementary Information accompanying this manuscript contains supplementary tables, robustness figures, and supporting material referenced in the text. Separately, the full reproducibility package, including code and curated datasets, is maintained in a private GitHub repository. Upon acceptance, the repository will be made publicly available, and the permanent public URL (and DOI, if archived) will be inserted here: [PUBLIC LINK/DOI TO BE ADDED UPON ACCEPTANCE]. No material-transfer agreements are required.

## CRediT authorship contribution statement

**Sabitoj Singh Virk:** Conceptualization; Methodology; Software; Data curation; Formal analysis; Investigation; Validation; Visualization; Writing – original draft; Writing – review and editing; Project administration. **Akashdeep Singh Virk:** Conceptualization; Methodology; Software; Data curation; Formal analysis; Investigation; Validation; Visualization; Writing – original draft; Writing – review and editing; Project administration.

