## Supplemental information for "Monoclonal Antibody Developability from Early Assay Panels: Machine Learning for Formulation and Pharmacokinetic Risk"

This document contains supplementary tables and robustness figures referenced in the manuscript. All curated datasets (DataS1-DataS5) and the full analysis code needed to reproduce the analyses and results reported in the manuscript and Supplementary Information are maintained as a single reproducibility package in a private GitHub repository; the package includes MANIFEST files enumerating all files and checksums. Upon acceptance, the repository will be made publicly available, and the permanent public URL (and DOI, if archived) will be inserted here: [PUBLIC LINK/DOI TO BE ADDED UPON ACCEPTANCE]. No material-transfer agreements are required.

**Table S1. Data dictionary for assay features and model outputs (variables appear exactly as column headers in the provided CSV files).**

| Task/Dataset | Variable | Description | Units | Notes / Source |
|---|---|---|---|---|
| Viscosity (Mock et al.) | DLS Interaction Parameter kD (mL/g) | Diffusion interaction parameter from DLS; proxy for net protein-protein interactions in dilute solution. | mL/g | Provided in source dataset. |
| Viscosity (Mock et al.) | SE-UHPLC Main Peak Plates (EP) | Theoretical plate count for the main SEC/SE-UHPLC monomer peak (peak sharpness/efficiency). | dimensionless (EP) | Peak-shape descriptor from SE-UHPLC. |
| Viscosity (Mock et al.) | AC-SINS Δλmax (nm) | AC-SINS spectral shift (Δλmax) relative to buffer/control; larger positive shifts indicate higher self-interaction. | nm | AC-SINS = affinity-capture self-interaction nanoparticle spectroscopy; Δλmax ≥ 15 nm is commonly used as a high self-interaction flag. |
| Viscosity (Mock et al.) | SE-UHPLC Main Peak FWHM (min) | Full width at half maximum of the main SEC/SE-UHPLC monomer peak (peak broadening). | min | Peak-shape descriptor from SE-UHPLC. |

| | | | | |
|---|---|---|---|---|
| Viscosity (Mock et al.) | Viscosity | Measured viscosity of concentrated antibody solution (prediction target). | cP | Provided as-is from source dataset. |
| Mouse PK (Liu et al.) | Heparin_RT | Retention time on heparin chromatography column. | min | Heparin chromatography is commonly used as a proxy for charge-driven nonspecific interactions. |
| Mouse PK (Liu et al.) | Heparin_pB_buffer | %B buffer at elution in heparin chromatography. | % | Represents elution strength/interactio n with heparin column. |
| Mouse PK (Liu et al.) | BVP_high | Baculovirus particle (BVP) binding signal (high condition). | unitless | Proxy for nonspecific interactions. |
| Mouse PK (Liu et al.) | poly_D_lysine | Binding signal to poly-D-lysine surface. | unitless | Proxy for charge-driven nonspecific interactions. |
| Mouse PK (Liu et al.) | AUCt | Area under the concentration-time curve to the last sampled timepoint (systemic exposure after IV dosing). | $10^6$ ng·h/mL | Higher AUCt generally corresponds to slower clearance. Values are reported as AUC0-672 h divided by $10^6$ (multiply by $10^6$to recover ng·h/mL). |
| Clinical outcome (Jain et al.) | AS | Accelerated stability (AS) SEC slope: long-term stability slope calculated from % aggregated measured by SEC during a 30-day incubation at 40 °C. | %/day | Defined in Jain et al. as “AS SEC slope” computed from % aggregated (SEC) vs time during accelerated stability testing. |
| Clinical outcome (Jain et al.) | PSR | Polyspecificity reagent binding assay. | unitless | Higher values indicate increased polyspecific interactions. |
| Clinical outcome (Jain et al.) | ACSINS | AC-SINS wavelength shift / readout. | nm | Self-interaction assay. |

| | | | | |
|---|---|---|---|---|
| Clinical outcome (Jain et al.) | ELISA | Multi-antigen ELISA readout. | fold-over-background (unitless) | Polyspecificity-related readout. |
| Clinical outcome (Jain et al.) | BVP | Baculovirus particle binding assay. | fold-over-background (unitless) | Polyspecificity-related readout. |
| Clinical outcome (Jain et al.) | Outcome | Internal-cohort binary label used for training/testing. | Approved / Terminated | Derived from publicly available status annotations as described in the manuscript. |
| Clinical outcome (external cohort) | Updated.Status | External-cohort label used for independent validation. | Approved / Terminated | As provided in the external cohort file. |
| HIC (Bailly et al.) | titer_mg_L | Expression titer / concentration. | mg/L | Bailly et al. developability panel. |
| HIC (Bailly et al.) | up_sec_pct_hmw | SEC % high-molecular-weight species (HMW). | % | Upstream SEC readout. |
| HIC (Bailly et al.) | up_sec_pct_main | SEC % main monomer peak. | % | Upstream SEC readout. |
| HIC (Bailly et al.) | up_sec_pct_lmw | SEC % low-molecular-weight species (LMW). | % | Upstream SEC readout. |
| HIC (Bailly et al.) | up_sec_retention_time_min | SEC retention time of main peak. | min | Upstream SEC readout. |
| HIC (Bailly et al.) | hp_rp_pct_pre | HP-RP chromatography: % pre-peak fraction. | % | Chromatography peak-area fraction. |
| HIC (Bailly et al.) | hp_rp_pct_main | HP-RP chromatography: % main-peak fraction. | % | Chromatography peak-area fraction. |
| HIC (Bailly et al.) | hp_rp_pct_post | HP-RP chromatography: % post-peak fraction. | % | Chromatography peak-area fraction. |
| HIC (Bailly et al.) | ce_sds_nr_pct_lmw | CE-SDS (non-reduced): % low-molecular-weight species. | % | CE-SDS NR readout. |
| HIC (Bailly et al.) | ce_sds_nr_pct_main | CE-SDS (non-reduced): % main species. | % | CE-SDS NR readout. |

| | | | | |
|---|---|---|---|---|
| HIC (Bailly et al.) | ce_sds_nr_pct_hmw | CE-SDS (non-reduced): % high-molecular-weight species. | % | CE-SDS NR readout. |
| HIC (Bailly et al.) | ce_sds_red_pct_lc | CE-SDS (reduced): % light chain (LC). | % | CE-SDS reduced readout. |
| HIC (Bailly et al.) | ce_sds_red_pct_hc | CE-SDS (reduced): % heavy chain (HC). | % | CE-SDS reduced readout. |
| HIC (Bailly et al.) | ce_sds_red_pct_others | CE-SDS (reduced): % other species. | % | CE-SDS reduced readout. |
| HIC (Bailly et al.) | nano_dsf_tonset_c | nanoDSF thermal onset temperature (Tonset). | °C | nanoDSF stability readout. |
| HIC (Bailly et al.) | nano_dsf_tm1_c | nanoDSF first melting temperature (Tm1). | °C | nanoDSF stability readout. |
| HIC (Bailly et al.) | nano_dsf_tagg_c | nanoDSF aggregation temperature (Tagg). | °C | nanoDSF stability readout. |
| HIC (Bailly et al.) | ac_sins_naac_ph_5_5 | AC-SINS λmax (maximum absorbance wavelength) measured in NaOAc buffer (pH 5.5). | nm | AC-SINS self-interaction assay readout reported as λmax in nm. |
| HIC (Bailly et al.) | ac_sins_pbs_ph_7_4 | AC-SINS λmax (maximum absorbance wavelength) measured in PBS buffer (pH 7.4). | nm | AC-SINS self-interaction assay readout reported as λmax in nm. |
| HIC (Bailly et al.) | low_ph_hold_up_sec_pct_hmw | Post low-pH hold: SEC % HMW. | % | Low-pH stress + SEC. |
| HIC (Bailly et al.) | low_ph_hold_up_sec_pct_main | Post low-pH hold: SEC % main peak. | % | Low-pH stress + SEC. |
| HIC (Bailly et al.) | low_ph_hold_up_sec_pct_lmw | Post low-pH hold: SEC % LMW. | % | Low-pH stress + SEC. |
| HIC (Bailly et al.) | low_ph_hold_up_sec_retention_time_min | Post low-pH hold: SEC retention time. | min | Low-pH stress + SEC. |
| HIC (Bailly et al.) | cief_pi | Isoelectric point (pI) from cIEF. | pH units | cIEF readout. |

| | | | | |
|---|---|---|---|---|
| HIC (Bailly et al.) | patch_cdr_hyd | Ensemble-average sum of hydrophobic patch surface area proximal to the CDR (structure-based). | Å$^2$ | Structure-derived surface patch area descriptor computed from Fab homology-model ensembles (reported in Bailly et al. as patch surface area, Å$^2$). |
| HIC (Bailly et al.) | patch_hyd | Ensemble-average sum of hydrophobic patch surface area over the Fab (structure-based). | Å$^2$ | Structure-derived surface patch area descriptor computed from Fab homology-model ensembles (reported in Bailly et al. as patch surface area, Å$^2$). |
| HIC (Bailly et al.) | patch_cdr_ion | Ensemble-average sum of ionic patch surface area proximal to the CDR (structure-based). | Å$^2$ | Structure-derived surface patch area descriptor computed from Fab homology-model ensembles (reported in Bailly et al. as patch surface area, Å$^2$). |
| HIC (Bailly et al.) | patch_ion | Ensemble-average sum of ionic patch surface area over the Fab (structure-based). | Å$^2$ | Structure-derived surface patch area descriptor computed from Fab homology-model ensembles (reported in Bailly et al. as patch surface area, Å$^2$). |
| HIC (Bailly et al.) | isotype_igg4 | Binary indicator for IgG4 isotype (1=IgG4, 0=non-IgG4). | binary | Isotype covariate. |
| HIC (Bailly et al.) | hic_rt_min | Hydrophobic interaction chromatography retention time (developability proxy). | min | Experimental HIC retention time; antibodies that did not elute were set to the maximum of 50 min in the source study. |

**Table S2. Bagged out-of-fold (OOF) ensemble performance for HIC retention time prediction on the Bailly et al. dataset (n = 152) using 10× repeated 5-fold cross-validation.**

| Model / feature set | $R^2$ (sklearn) | Pearson $r^2$ | Spearman ρ | RMSE (min) | MAE (min) | Balanced Acc@30 | MCC@30 |
|---|---|---|---|---|---|---|---|
| ACeT (assays only) | 0.796 | 0.825 | 0.881 | 4.995 | 3.256 | 0.891 | 0.777 |
| ACeT (assays + patch) | 0.756 | 0.786 | 0.875 | 5.459 | 3.365 | 0.895 | 0.787 |
| ACeT (patch only) | 0.545 | 0.585 | 0.677 | 7.451 | 5.019 | 0.823 | 0.652 |
| Bailly baseline (patch linear, refit) | 0.506 | 0.507 | 0.652 | 7.763 | 6.444 | 0.777 | 0.550 |

*Notes:* For each CV replicate (one complete 5-fold run), every antibody receives an OOF prediction from a model that did not train on that antibody. The reported "bagged OOF" prediction for each antibody is the average of its OOF predictions across the 10 CV replicates, and metrics are computed once on these 152 averaged OOF predictions. Pearson $r^2$ is the squared Pearson correlation (as reported in the original study); $R^2$ (sklearn) refers to the coefficient of determination. Binary triage at 30 min is computed by thresholding predicted HIC RT: predicted "good" if pred ≤ 30 min, predicted "poor" if pred > 30 min (true labels: good ≤ 30, poor > 30). Balanced Acc@30 = (good_acc + poor_acc)/2. MCC@30 is the Matthews correlation coefficient computed from the 2×2 confusion matrix at the same 30-min cutoff: (TP·TN − FP·FN) / √((TP+FP)(TP+FN)(TN+FP)(TN+FN)).

**Table S3. CV-replicate variability of out-of-fold (OOF) performance for HIC retention time prediction (10× repeated 5-fold CV; mean ± SD across CV replicates).**

| Model / feature set | Pearson $r^2$ (mean ± SD) | Spearman ρ (mean ± SD) | RMSE (min, mean ± SD) | MAE (min, mean ± SD) | Balanced Acc@30 (mean ± SD) | MCC@30 (mean ± SD) |
|---|---|---|---|---|---|---|
| ACeT (assays only) | 0.725 ± 0.029 | 0.852 ± 0.017 | 6.669 ± 0.753 | 3.635 ± 0.532 | 0.887 ± 0.029 | 0.773 ± 0.057 |
| ACeT (assays + patch) | 0.695 ± 0.037 | 0.842 ± 0.022 | 6.886 ± 0.635 | 3.812 ± 0.379 | 0.885 ± 0.021 | 0.769 ± 0.042 |

| | | | | | | |
|---|---|---|---|---|---|---|
| ACeT (patch only) | 0.540 ± 0.038 | 0.665 ± 0.022 | 8.064 ± 0.441 | 5.129 ± 0.341 | 0.820 ± 0.017 | 0.644 ± 0.036 |
| Bailly baseline (patch linear, refit) | 0.504 ± 0.015 | 0.649 ± 0.007 | 7.789 ± 0.124 | 6.457 ± 0.110 | 0.771 ± 0.007 | 0.540 ± 0.012 |

*Notes:* Pearson $r^2$ is the squared Pearson correlation; $R^2$ refers to the coefficient of determination. Binary triage uses cutoff = 30 min (good ≤ 30, poor > 30). Metrics were computed separately within each CV replicate using that replicate's OOF predictions (n = 152 per replicate), then summarized as mean ± SD across the 10 CV replicates, reflecting sensitivity to fold assignment / training stochasticity. MCC@30 is the Matthews correlation coefficient at the 30-min cutoff (computed from TP, TN, FP, FN as above).

**Table S4. Simple single-descriptor baselines for viscosity prediction on the seed-0 train/test split (n = 52/23).**

| Model | Input features | Preprocessing | Test $R^2$ | Test RMSE (cP) | Test MAE (cP) |
|---|---|---|---|---|---|
| Ridge (kD-only) | DLS interaction parameter kD | Quantile Transformer (normal) + Ridge (α=1.0) | 0.27 | 8.02 | 5.89 |
| Ridge (kD + SEC plates) | kD; SE-UHPLC main peak plates (EP) | Quantile Transformer (normal) + Ridge (α=1.0) | 0.48 | 6.76 | 5.16 |
| ACeT (four assays; manuscript Fig. 2) | kD; SEC plates; SEC FWHM; AC-SINS Δλmax | Full ACeT pipeline (see Methods) | 0.75 | 4.8 | 3.0 |

*Notes*: Ridge baselines are simple regressions intended to contextualize the value of adding SEC peak-shape descriptors beyond kD alone. ACeT metrics are reported from the main manuscript (Fig. 2) for comparison.

### Supplementary Figures

Key robustness figures for the three endpoints reported in the manuscript. Robustness was evaluated by repeating model training/evaluation across multiple random train/test partitions (different random seeds) and aggregating out-of-sample behavior.

**Figure S1. Viscosity robustness: pooled out-of-sample absolute log10 error CDF across split seeds 0-5.**

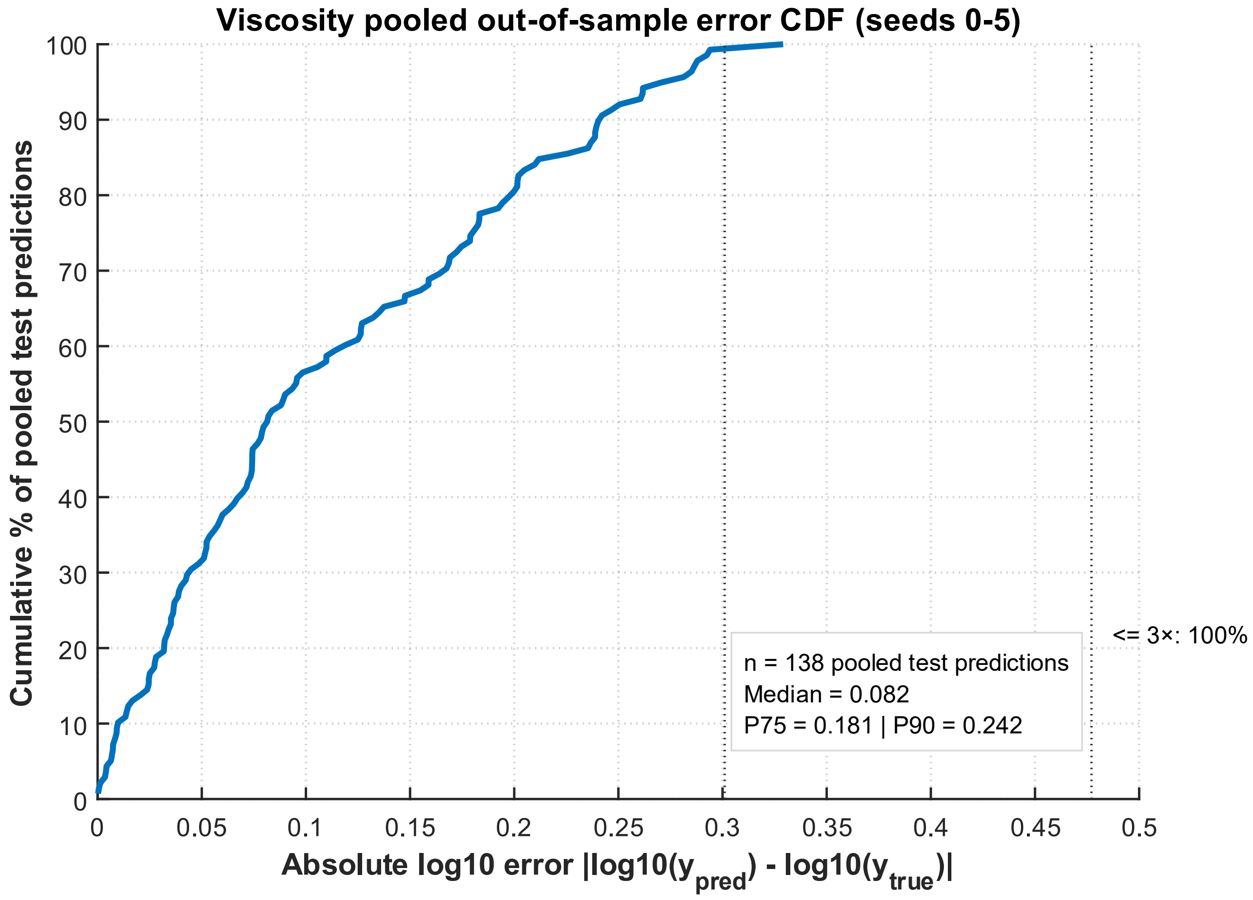

**Figure S2. Viscosity robustness: out-of-sample parity across composite splits (log-log) for split seeds 0-5 (seed 0 corresponds to the manuscript split).**

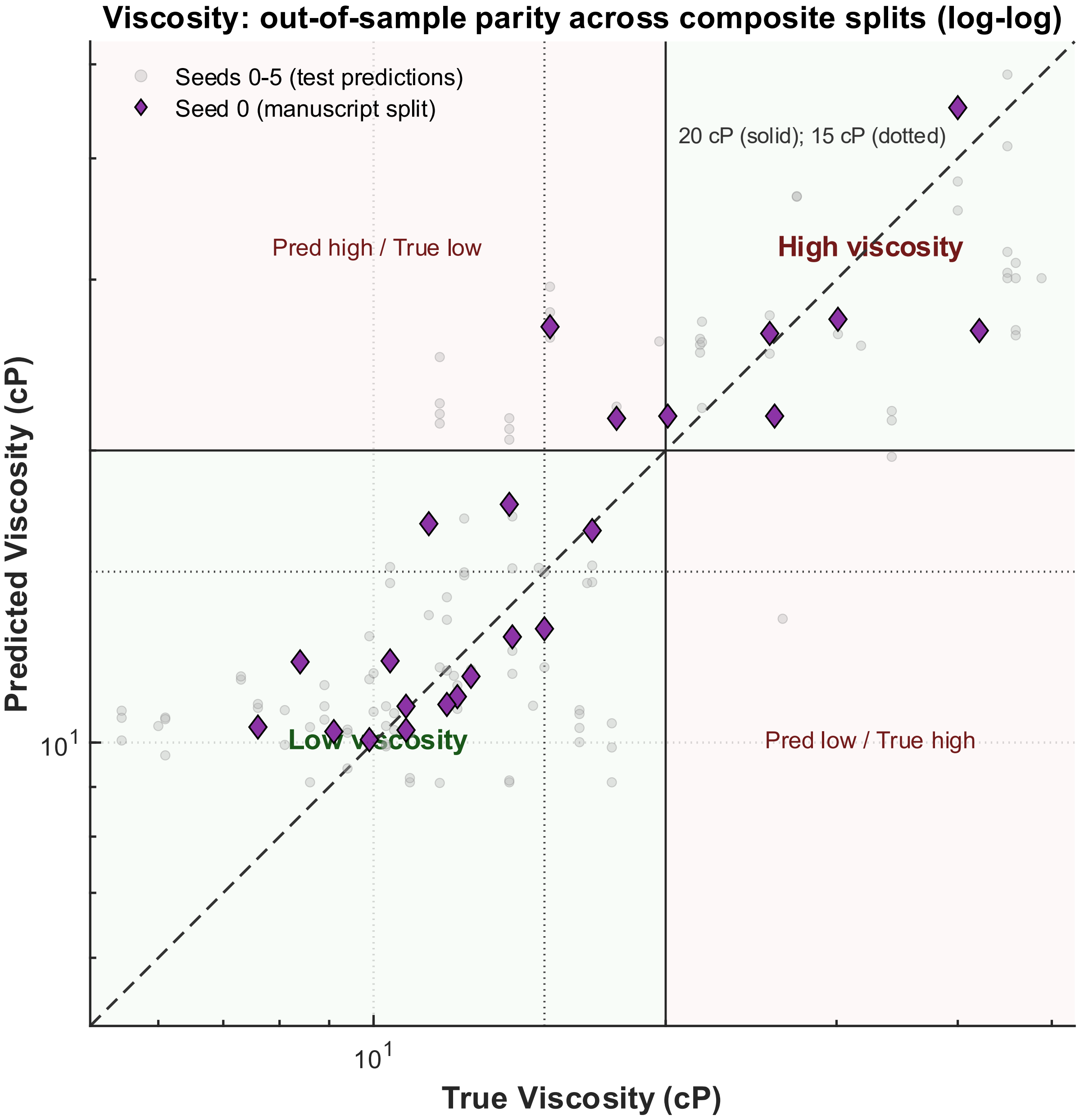

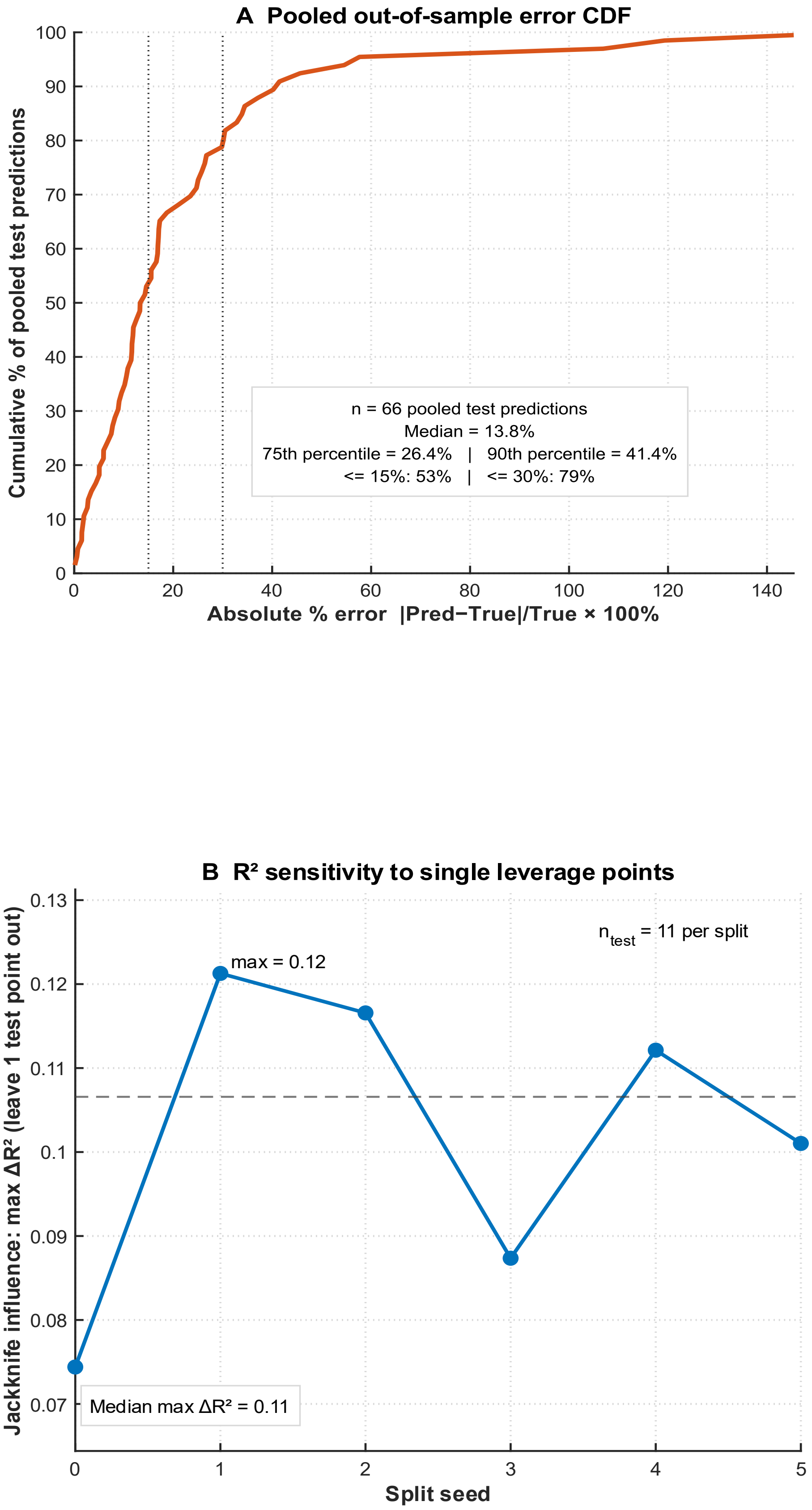


**Figure S3. Mouse IV clearance proxy robustness: pooled out-of-sample absolute percent error CDF across split seeds 0-5.**

**Figure S4. Mouse IV clearance proxy robustness: out-of-sample parity across composite splits (log–log) for split seeds 0-5 (seed 0 corresponds to the manuscript split).**

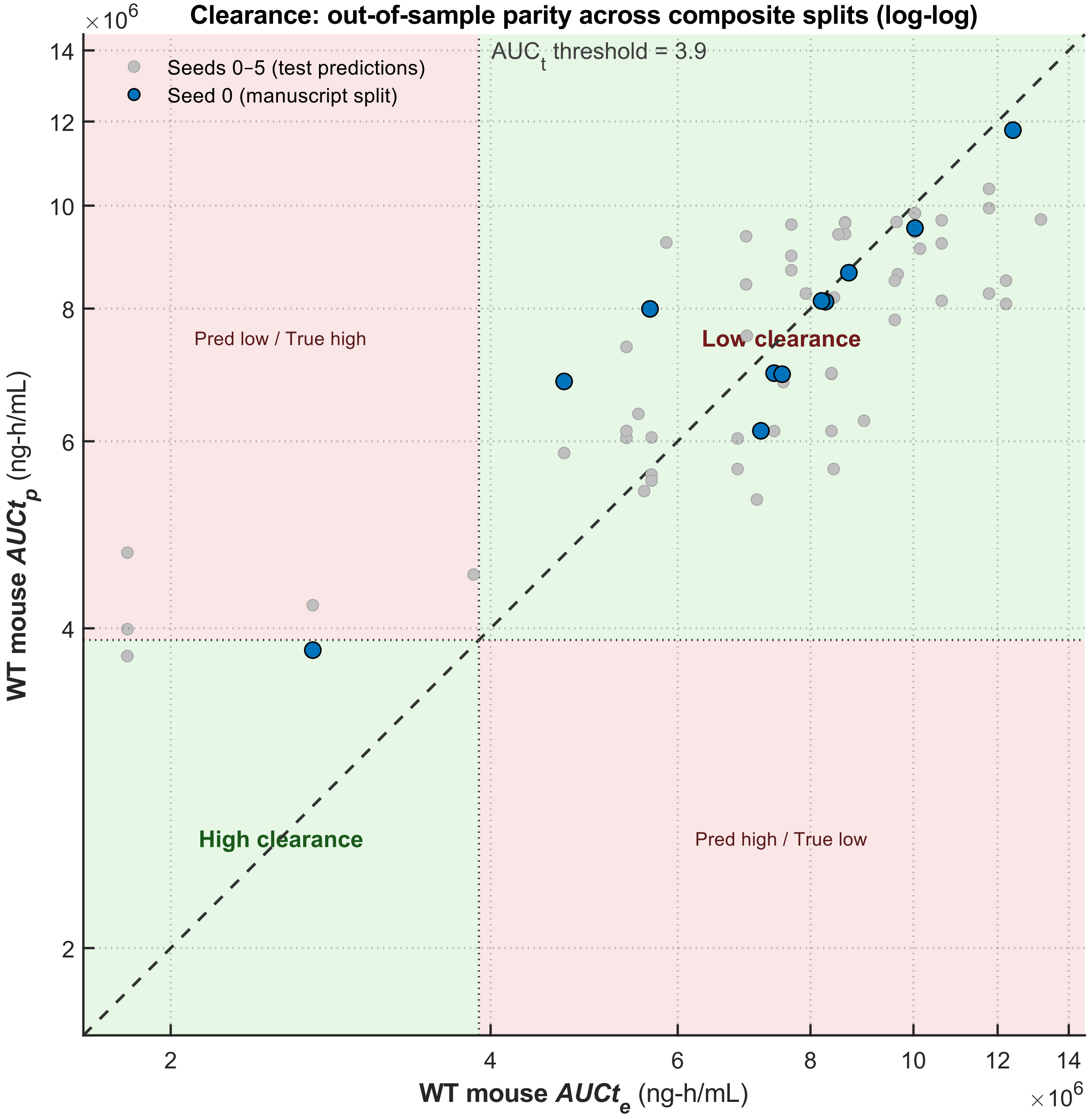

**Figure S5. Clinical outcome robustness (external evaluation): external cohort balanced accuracy, pooled log-loss distribution, and jackknife sensitivity across split seeds 0-10. Terminated denotes the negative class (by the May 2025 cutoff).**

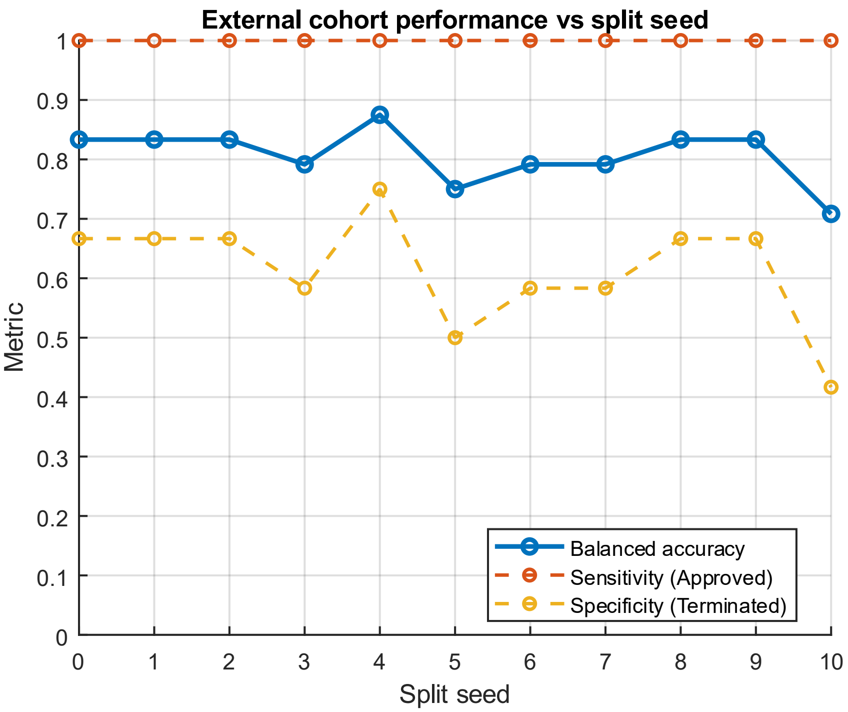


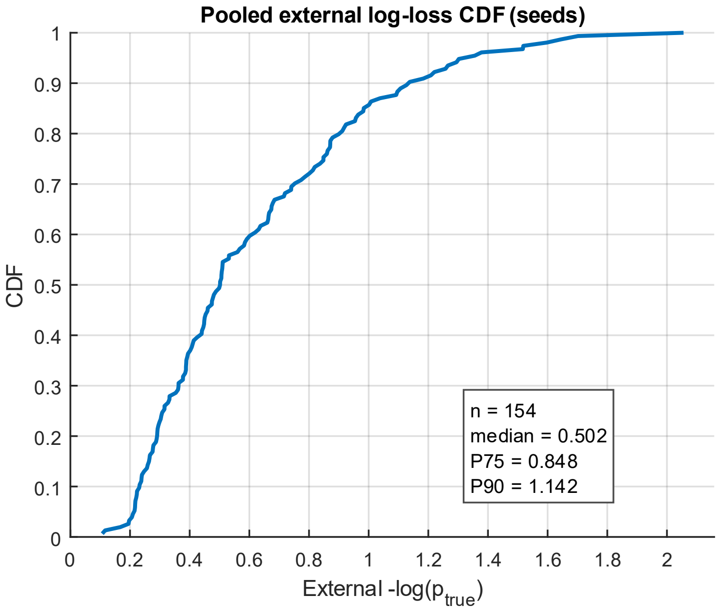


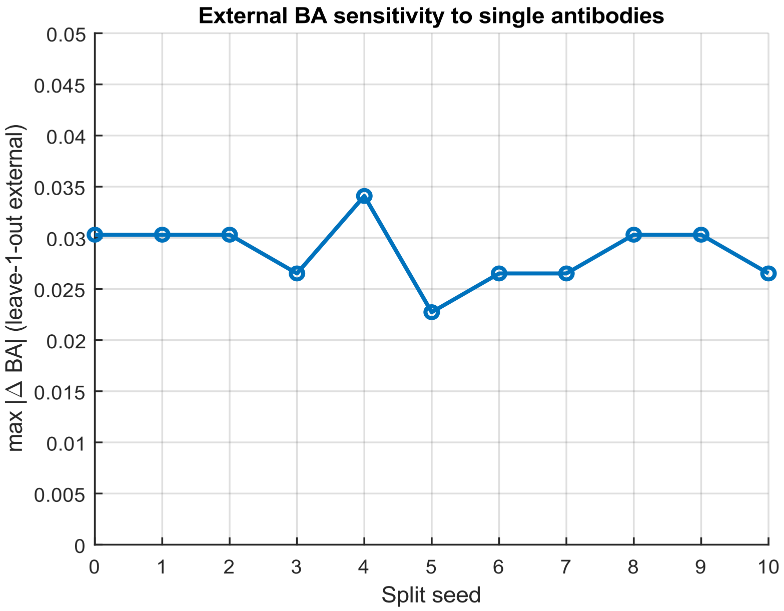

**Figure S6. HIC retention time prediction (Bailly et al.): out-of-fold (OOF) parity for ACeT using assays-only inputs. Predictions are bagged OOF values averaged across 10 repeats of 5-fold CV; dashed line indicates unity.**

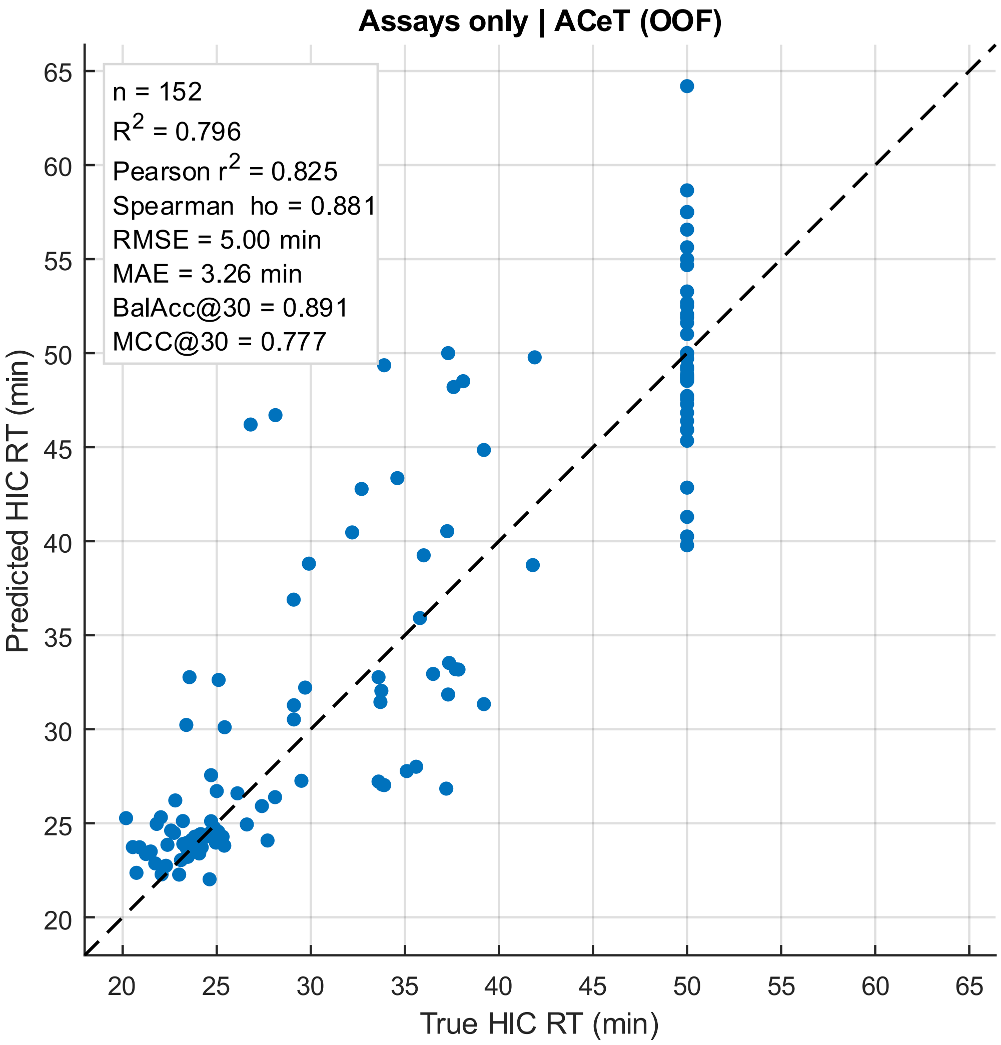

**Figure S7. HIC retention time prediction (Bailly et al.): out-of-fold (OOF) parity for ACeT using patch descriptors only (apples-to-apples vs patch-only baseline). Predictions are bagged OOF values averaged across 10 repeats of 5-fold CV; dashed line indicates unity.**

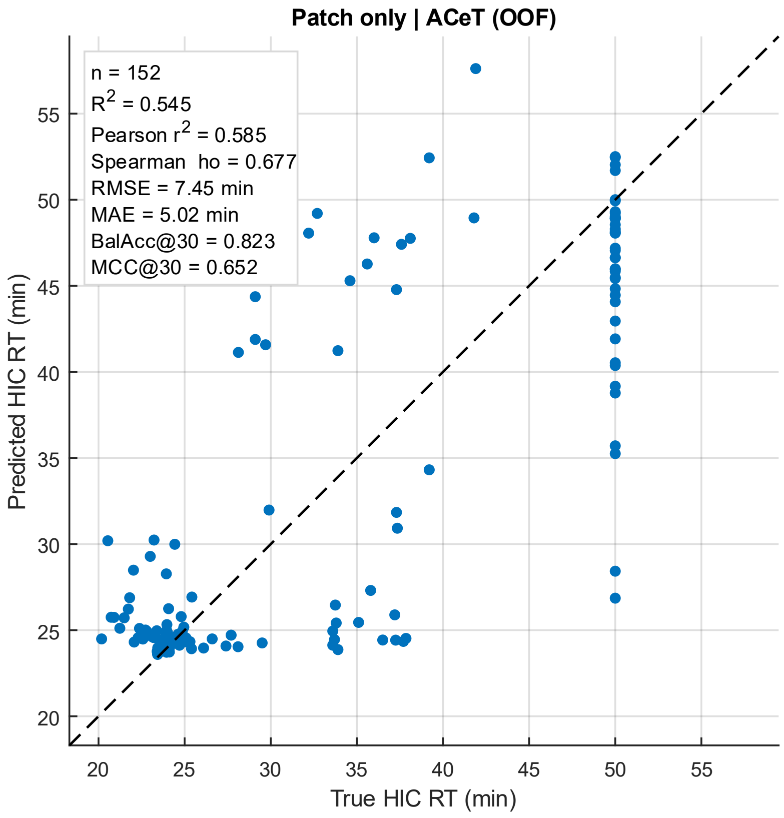


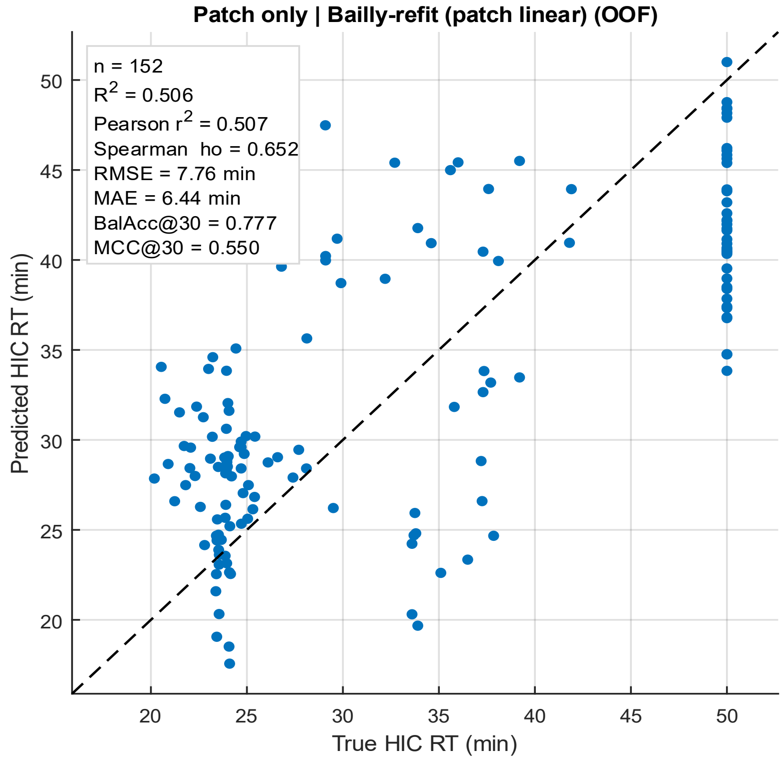

**Figure S8. Binary developability triage at 30 min on the Bailly et al. HIC dataset using bagged OOF predictions (10× repeated 5-fold CV).** Confusion matrices compare **ACeT (assays-only)** versus the fold-matched **Bailly-refit baseline** when converting continuous HIC RT predictions into a binary decision at the **30-min cutoff** (true label: *good* ≤ 30 min, *poor* > 30 min; predicted label thresholded identically on predicted HIC RT). Counts are computed on the **bagged OOF predictions** (per-antibody OOF predictions averaged across the 10 CV replicates). Balanced accuracy and MCC corresponding to this triage rule are reported in **Table S2**, and replicate-level variability in **Table S3**.

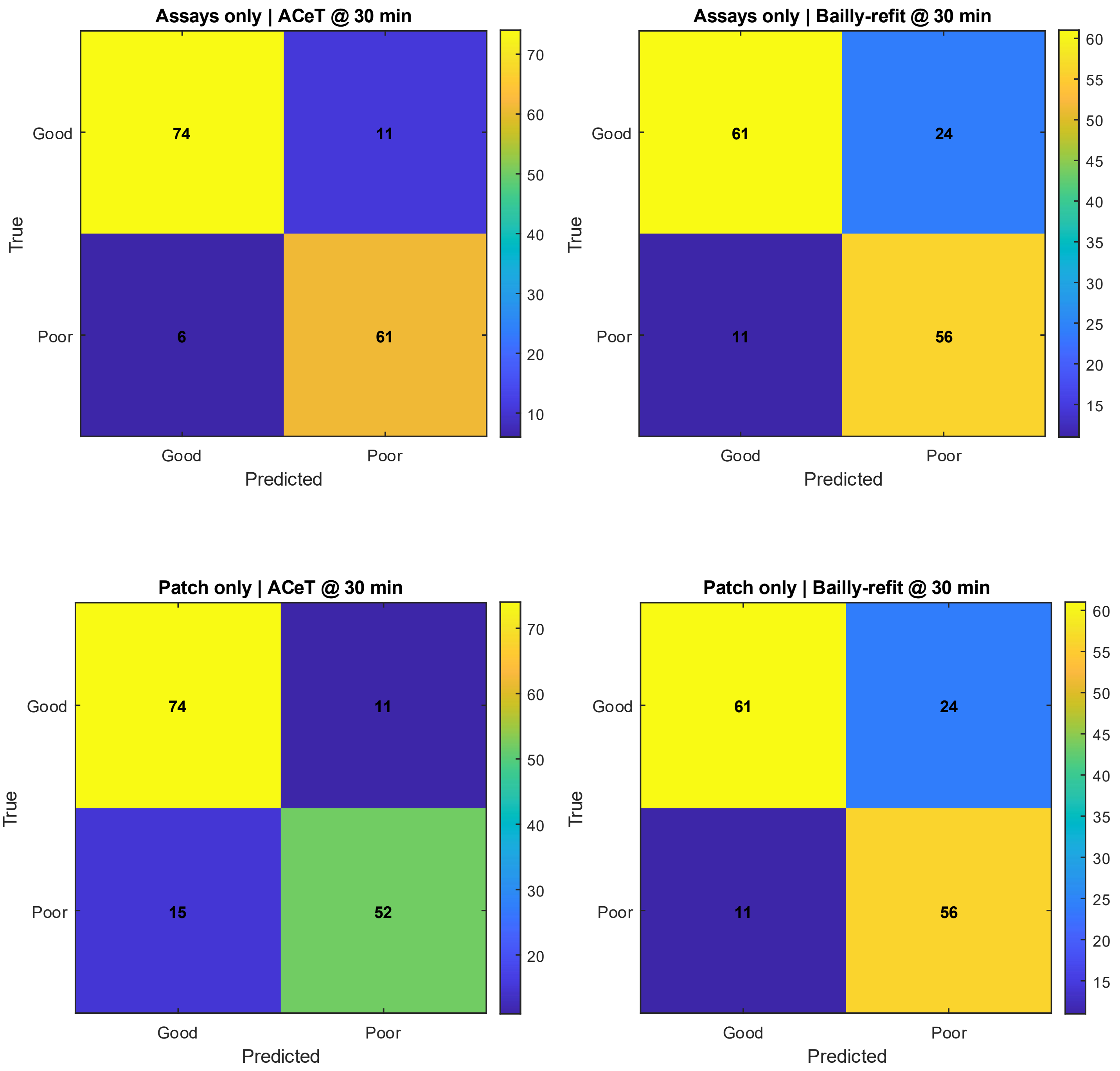

**Figure S9. Relationship between SEC monomer peak-shape metrics and high-concentration viscosity in the 75-antibody viscosity dataset (DataS1). Left: viscosity versus SE-UHPLC main peak plates (EP). Right: viscosity versus SE-UHPLC main peak FWHM (min). Broad peaks (lower plate count and higher FWHM) are associated with higher viscosity, consistent with the interpretation that SEC peak dispersion can reflect heterogeneity, reversible self-association, and/or secondary interactions.**

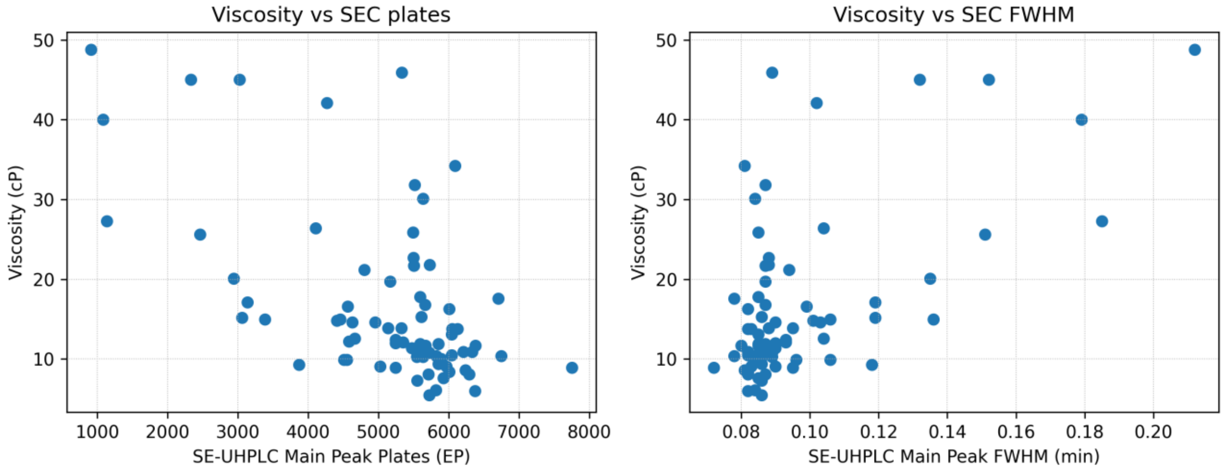
